# The *Caenorhabditis elegans* TDRD5/7-like protein, LOTR-1, interacts with the helicase ZNFX-1 to balance epigenetic signals in the germline

**DOI:** 10.1101/2021.06.18.448978

**Authors:** Elisabeth A. Marnik, Miguel V. Almeida, P. Giselle Cipriani, George Chung, Edoardo Caspani, Emil Karaulanov, Falk Butter, Catherine S. Sharp, John Zinno, Hin Hark Gan, Fabio Piano, René F Ketting, Kristin C. Gunsalus, Dustin L. Updike

## Abstract

LOTUS and Tudor domain containing proteins have critical roles in the germline. Proteins that contain these domains, such as Tejas/Tapas in *Drosophila*, help localize Vasa to the germ granules and facilitate piRNA-mediated transposon silencing. The homologous proteins in mammals, TDRD5 and TDRD7, are required during spermiogenesis. Until now, proteins containing both LOTUS and Tudor domains in *Caenorhabditis elegans* have remained elusive. Here we describe LOTR-1 (D1081.7), which derives its name from its **<u>LO</u>**TUS and **<u>T</u>**udo**<u>r</u>** domains. Interestingly, LOTR-1 docks next to P granules to colocalize with the broadly conserved Z-granule helicase, ZNFX-1. LOTR-1’s Z-granule association requires its Tudor domain, but both LOTUS and Tudor deletions affect brood size when coupled with a knockdown of the Vasa homolog *glh-1*. In addition to interacting with the germ-granule components WAGO-1, PRG-1 and DEPS-1, we identified a Tudor-dependent association with ZNFX-1. Like *znfx-1* mutants, *lotr-1* mutants lose small RNAs from the 3’ ends of WAGO and Mutator targets, reminiscent of the loss of piRNAs from the 3’ ends of piRNA precursor transcripts in mouse Tdrd5 mutants. Our work suggests that LOTR-1 acts in a conserved mechanism that brings small RNA generating mechanisms towards the 3’ ends of small RNA templates or precursors.

## Introduction

Germ cells produce the next generation, and their pluripotent potential is instrumental for ensuring fertility and proper development. While germline and somatic cells contain the same DNA, differences within their cytoplasm, or germplasm, help distinguish their respective fates.[1] In some animals, ectopic germplasm can be sufficient to reprogram somatic nuclei.[2–5] Additionally, the presence of germ plasm components in the soma promotes cell proliferation, pluripotency, and tumorigenesis.[6–9] Understanding how the germplasm derives this reprogramming potential is a critical undertaking with far-reaching implications to reproductive, developmental, and regenerative biology.

The germplasm contains non-membrane-bound ribonucleoprotein (RNP) condensates called germ granules that harbor part of this reprogramming potential.[10,11] One conserved feature of germ granules across species is the presence of proteins with LOTUS and Tudor domains. LOTUS is a name derived from the germ-granule-associated proteins **<u>L</u>**imkain/MARF1, **<u>O</u>**skar, and **<u>Tu</u>**dor-containing protein**<u>s</u>** 5 and 7.[12–15] The LOTUS domain takes on a winged helix-turn-helix (wHTH) folding pattern to facilitate RNA and protein interactions critical to germline development. *Drosophila* Oskar uses its LOTUS domain to self-dimerize and interface with the Vasa DEAD-box helicase, stimulating its activity to promote piRNA amplification in the germline.[16] Oskar expression in the *Drosophila* oocyte drives germ-granule assembly, while its mislocalization is sufficient to form ectopic germ cells from somatic progenitors.[17,18] However, drawing parallels between Oskar and other LOTUS-containing proteins is difficult as Oskar is confined to some insects and likely arose *de novo via* fusion of a eukaryotic LOTUS domain with a bacterially derived hydrolase-like domain called OSK through horizontal gene transfer.[19] In *Drosophila*, Limkain/MARF-1 regulates oocyte maturation and its LOTUS domain associates with the CCR4-NOT deadenylase complex. This results in the shortening of poly-A tails and translational regulation of targeted RNA transcripts.[20] In mice, MARF1 localizes to germ granules in oocytes, where it is critical for normal oocyte development. Female mice lacking MARF1 are sterile due to failures in oocyte meiosis and increased retrotransposon activity.[21,22] In addition to interfacing with proteins, LOTUS domains also bind RNA with a preference for G-rich RNAs and those that form a G-quadruplex (G4) secondary structure.[23] The function of these G-rich and G4 interactions remains unclear but could be instrumental to the role of LOTUS in RNA metabolism and regulation.[23]

TDRD5 and TDRD7 contain LOTUS domains paired with Tudor scaffolding domains.[24] Tudor domains have been shown to bind methylated arginines and lysines, with some preference for Argonaute proteins and histone tails.[25–27] In mammals, TDRD5 and TDRD7 associate with key components of the piRNA pathway and are required for normal spermatogenesis and retrotransposon silencing.[28–32] The TDRD5 and TDRD7 orthologs in *Drosophila*, respectively known as Tejas and Tapas, are required for proper germ granule formation and piRNA silencing of transposons in the germline.[33,34] Combined, these findings illustrate the importance of germ-granule localized LOTUS and Tudor domain-containing proteins in maintaining germline integrity through translational regulation, transposon silencing, and stimulation of Vasa helicase activity.

Germ granule studies are aided in *C. elegans* by the nematode’s transparency, permitting the observation of germ granules (or P granules) in living animals at all stages of development. During embryonic development, P granules segregate to germline blastomeres (P cells) before coming to reside in two primordial germ cells (PGCs).[35] Following the formation of PGCs, distinct sub-granules emerge from P granules to occupy neighboring sites at the nuclear periphery.[36] Known sub-granules include Z-granules, SIMR foci, and Mutator foci -each containing sets of proteins that refine and resolve processes or steps of small RNA biosynthesis.[37– 39]

*C. elegans* expresses several classes of small RNAs.[40,41] 21U-RNAs are considered the piRNAs of *C. elegans* due to their interaction with the germline-expressed PRG-1 Argonaute, the main Piwi-class Argonaute of *C. elegans*.[42–44] Similar to piRNAs of other organisms, PRG-1/21U-RNAs target “non-self” sequences such as transposable elements (TEs).[42,45,46] Target recognition leads to the recruitment of RNA-dependent RNA Polymerases (RdRPs), which synthesize complementary 22G-RNAs from template target RNAs.[45,46] In turn, 22G-RNAs associate with WAGO-class Argonautes, which elicit target gene silencing at the post-transcriptional and transcriptional levels.[47–51] An important function of PRG-1/21U-RNAs is to prevent the erroneous targeting of essential genes by 22G-RNAs.[52,53] Gene silencing initiated by 21U-RNAs can become independent of the initial PRG-1/21U-RNA trigger and self-sustained by 22G-RNAs and heterochromatin marks in ways that are not fully understood.[54–56] This PRG-1/21U-RNA-independent silencing is termed RNA-induced epigenetic silencing (RNAe).

26G-RNAs are produced by the RdRP RRF-3 and additional cofactors in gonads.[57–62] Two distinct subpopulations of 26G-RNAs are expressed: those expressed during spermatogenesis that associate with the Argonautes ALG-3/4[59,63,64], and those expressed during oogenesis and embryogenesis that associate with the Argonaute ERGO-1.[57,59,65] 26G-RNAs also trigger secondary 22G-RNA synthesis to produce robust target gene silencing. Due to their production downstream of several primary pathways, 22G-RNAs comprise a highly complex small RNA species. 22G-RNAs can be functionally divided into distinct subpopulations based on their expression pattern, the WAGO-class Argonaute protein with which they interact, and their set of target genes.[40,41]

In *C. elegans*, aspects of small RNA biogenesis and gene silencing occur both in the nucleus and cytoplasm.[40] Cytoplasmic reactions of small RNA pathways mostly take place in germ granules, as extrapolated by the localization of many small RNA cofactors to these granules.[66] The interplay of small RNA biogenesis and silencing seems complex, but the partitioning of germ granules into sub-compartments suggests that these processes are physically organized in a vectorial manner, similar to the multiphase liquid condensates that mediate ribosome biogenesis in the nucleolus.[67] For example, Mutator foci are considered the site of WAGO-class 22G-RNA biogenesis.[68] Z granules were initially defined by the localization of ZNFX-1, an RNA helicase required for inheritance of small RNAs and transgenerational germ cell homeostasis.[69] Interestingly, ZNFX-1 was shown to interact with Argonaute proteins and is required for the correct positioning of RdRPs in their target transcripts.[69,70] *znfx-1* mutants display unbalanced 22G-RNA synthesis, with higher 22G-RNA levels produced towards the 5’ of the target transcript. PID-2/4/5 are recently identified factors that affect the structure of Z granules, are required for germ cell immortality, and are similarly required to balance 22G-RNA signals.[38,39] These studies demonstrate a link between Z granules and the biogenesis and inheritance of 22G-RNAs.

The role of LOTUS-domain proteins in *C. elegans* has remained unexplored. However, three LOTUS containing proteins have recently been identified: MIP-1, MIP-2, and D1081.7.[71] We find that D1081.7 is in germ granules and interacts with both MIP-1 and MIP-2. D1081.7 is the only known *C. elegans* protein to pair LOTUS and Tudor domains, similar to both TDRD5 and TDRD7, so we have named it **LO**TUS and **T**udo**R** containing protein **1** (LOTR-1). The Tudor domain of LOTR-1 is required for its association with germ-granules, but its LOTUS domains are not. Mass spectrometry revealed a robust reciprocal association between LOTR-1 and ZNFX-1, and imaging showed that LOTR-1 partitions with ZNFX-1 into Z granules with the formation of PGCs. Furthermore, LOTR-1, like ZNFX-1, is required for the normal distribution of PRG-1, MIP-1 and MIP-2 in the germline, but is largely dispensable for localization of the core P-granule components GLH-1, DEPS-1, PGL-1 and PGL-3 in adult germ cells. *lotr-1* mutants, including one with an in-frame LOTUS deletion, have deregulated 22G-and 26G-RNAs, and altered distribution of WAGO/Mutator class 22G-RNAs towards the 5’ end of some transcripts, similar to what has been described for *znfx-1* mutants.[70] These *lotr-*1 mutants also have a mortal germline phenotype, a common feature in small RNA pathway mutants. Therefore, we conclude that LOTR-1 functions with ZNFX-1 in Z granules to ensure balanced 22G-RNA biogenesis and the proper silencing of mutator targets from one generation to the next. These findings promise to shed light on both the somatic and germline functions of TDRD5, TDRD7, and other proteins containing paired LOTUS and Tudor domains.

## Results

### LOTR-1 is a germ-granule protein that contains both LOTUS and Tudor domains

Inaugural papers that first described the LOTUS winged-helix domain and its conservation identified *C. elegans* D1081.7 as a hypothetical orphan protein with a single LOTUS domain.[12,13] More recently, LOTUS domains have been subdivided into extended LOTUS (eLOTUS) domains (like the LOTUS domain in *Drosophila*’s Oskar), and minimal LOTUS (mLOTUS) domains that lack a conserved α5 helix (like those present in mammalian MARF1) (S1 Fig).[72] The LOTUS domain of D1081.7 described in these inaugural papers corresponds to the eLOTUS domain (aa 33-130), but our analysis uncovered an accompanying mLOTUS domain (aa 180-285) (Fig 1A). LOTUS domains have low sequence similarity and are challenging to identify using sequence analysis alone. The mLOTUS domain of D1081.7 has a strong sequence identity (21%) to the winged-helix region of Cdt1, a regulator in the DNA replication complex.[73] The predicted 3D structures of D1081.7 LOTUS domains superimpose well with solved structures from other species: eLOTUS domains of D1081.7 and Oskar align with a root-mean-square deviation (rmsd) of 2.8 Å, whereas mLOTUS, which lacks the α5 helix, deviates from both fly and human LOTUS folds by ∼3.5 Å rmsd (Fig 1B). Paired LOTUS domains reflect the arrangement of TDRD5 and TDRD7 in mammals and two proteins recently described in *C. elegans* called MIP-1 and MIP-2 (S1 Fig).[71]

**Figure 1.**
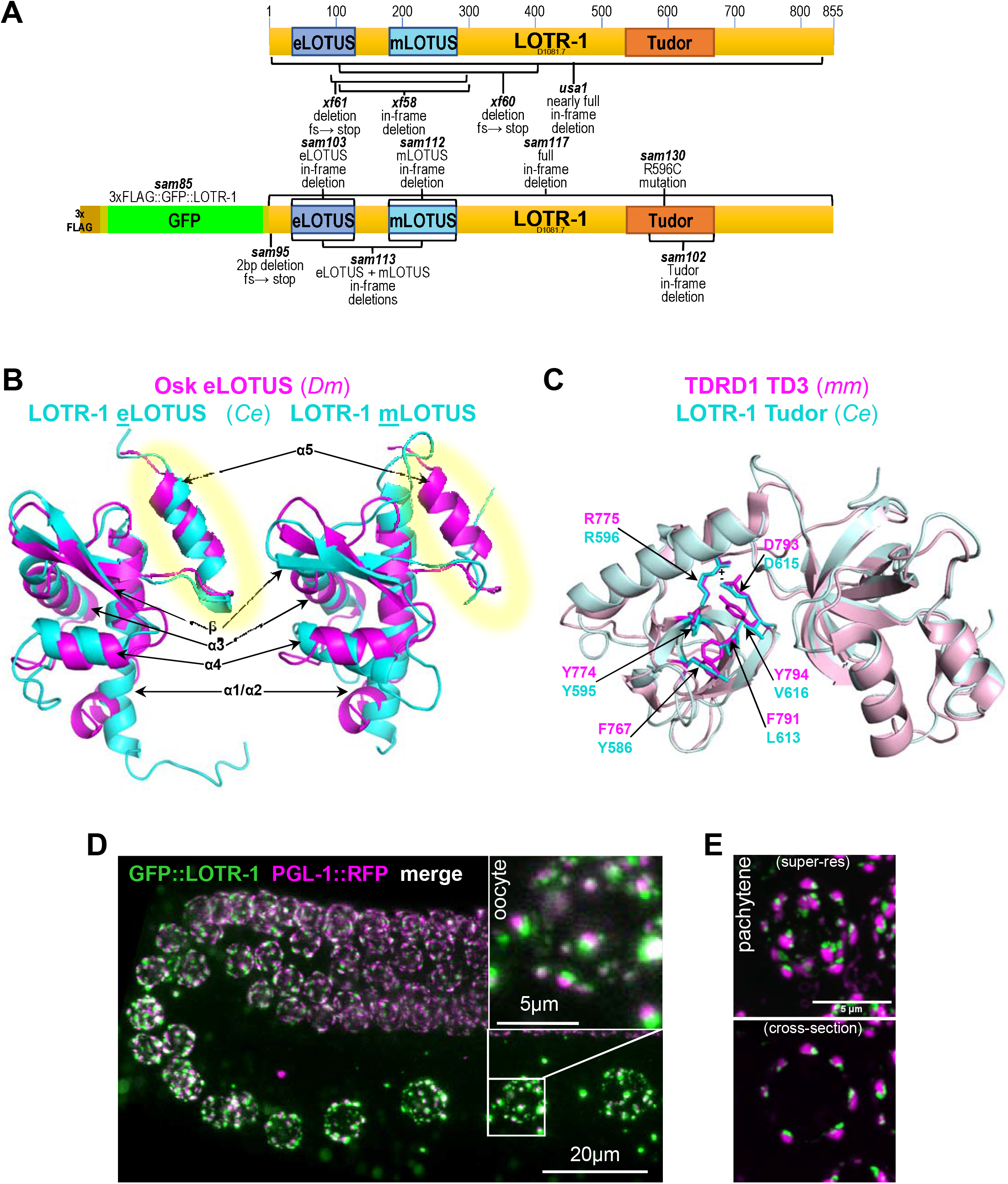
LOTR-1 is a germ granule protein that contains LOTUS and Tudor domains. A) Top schematic depicts the location of the eLOTUS, mLOTUS and Tudor domain of LOTR-1. Bottom depicts the CRISPR edit used to add an N terminal 3X FLAG tag and GFP. Alleles generated for this study are indicated. B) Predicted 3D model overlap of the eLOTUS and mLOTUS domains of *C. elegans* LOTR-1 with the Oskar eLOTUS domain from *Drosophila melanogaster*. The α5 helix (highlighted) is present in the eLOTUS domain. C) Predicted 3D structure of the Tudor domain (aa 534-670) of LOTR-1 overlapped with TD3 domain of mouse TDRD1. D) GFP::LOTR-1 and PGL-1::RFP in the germline of living worms. E) Super-resolution confocal imaging of GFP::LOTR-1 and PGL-1::RFP in pachytene germ cells.

Default BLAST parameters fail to uncover the Tudor domain within D1081.7, but it readily appears in domain-enhanced searches and was previously identified using multiple sequence alignment.[12] The Tudor domain (aa 534-670) is most similar to Tudor domain 3 (TD3) of mouse TDRD1, which is known to bind symmetrically dimethylated arginine (sDMA) marks through a hydrophobic pocket that is created by the arrangement of four aromatic residues (Fig 1C, F767, Y774, Y791, and Y794).[74] This pocket is stabilized by charge interactions between R775 and D793. The position of R596 and D615 is conserved in the Tudor domain of D1081.7, but the pocket replaces two of the four aromatic residues with other hydrophobic alternatives (L613 and V616). How this impacts sDMA binding is unknown.

The combination of both LOTUS and Tudor domains in D1081.7 is unique to homologs of Tejas/TDRD5 and Tapas/TDRD7 in *Drosophila* and mammals, which are piRNA pathway-regulating components that localize to the perinuclear nuage of germ cells. According to our BLAST searches, D1081.7 is the only protein in *C. elegans* known to have both LOTUS and Tudor domains. The Tudor and LOTUS domains of TDRD7 and TDRD5 are arranged similarly to those of D1081.7, with TDRD5 sharing 34% sequence similarity to D1081.7 across 466 aa, and TDRD7 sharing 39% sequence similarity across 550 aa. Its domain architecture and predicted structure suggests that D1081.7 is the homolog of TDRD5 and TDRD7 in *C. elegans*. Given these structural features, D1081.7 was named as <u>LO</u>TUS and <u>T</u>udor domain protein <u>1</u> (LOTR-1).

To determine the expression of LOTR-1, CRISPR/Cas9 genome editing was used to place an N-terminal GFP tag on endogenous *lotr-1* in a strain carrying PGL-1::RFP. LOTR-1 localized to germ granules throughout the adult germline; however, LOTR-1 granules appeared docked next to P granules marked by PGL-1::RFP (Fig 1D). Super-resolution confocal imaging of pachytene germ cells confirmed that LOTR-1 and PGL-1 granules appear adjacent to one another in the adult germline (Fig 1E). This pattern is similar to the P-granule docking of Z granules, Mutator foci, and SIMR foci, suggesting that LOTR-1 could reside within these germ granule sub-compartments.[69,70,75]

### Functional analysis of LOTR-1 domains reveals effects on subcellular localization and fertility

To understand how LOTR-1 is recruited to germ granules, a series of point mutations and deletions was introduced into the 3xFLAG::GFP::LOTR-1 strain (Fig 1A). Deletions of the mLOTUS and combined eLOTUS/mLOTUS domains fail to disperse truncated LOTR-1 from germ granules in young adults (Fig 2A). In contrast, the Tudor domain deletion causes LOTR-1 to disperse from most germ granules in the adult germline. A point mutation in the conserved arginine (R596C) of LOTR-1, which is predicted to stabilize the pocket with the potential to bind sDMAs, disperses LOTR-1 to the same degree as the deletion of the Tudor domain (Fig 2A).[37] These results suggest that the germ-granule association of LOTR-1 depends primarily on its Tudor domain, potentially mediated through well-characterized Tudor/sDMA interactions. Interestingly, in the absence of the Tudor domain, LOTR-1 is still retained in germ granules during spermatogenesis in the fourth larval stage (Fig 2B, orange box), suggesting that its retention in spermatogenic germ granules could be independent of Tudor/sDMA interactions.

**Figure 2.**
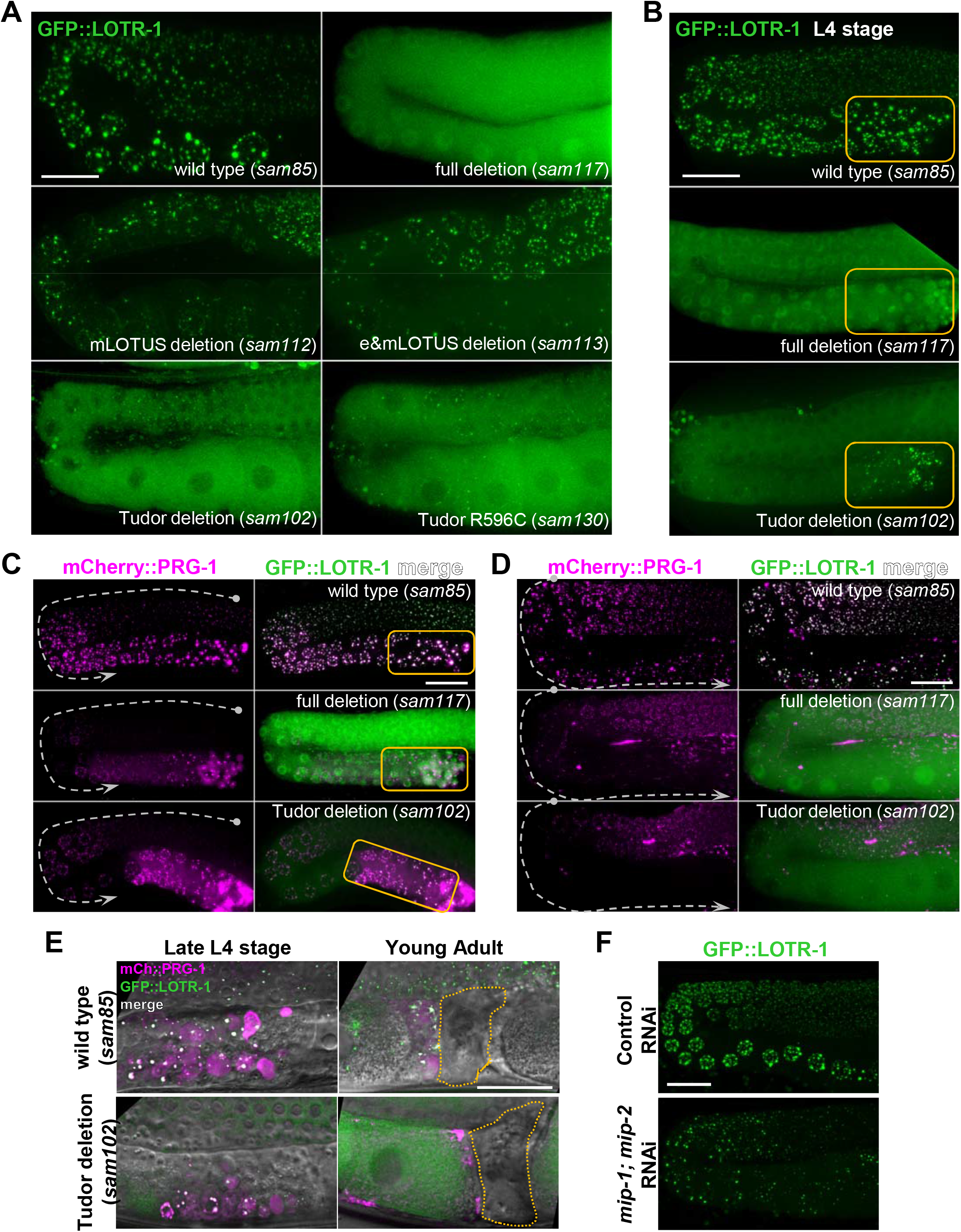
The germ granule localization of LOTR-1 is dependent on its Tudor domain and the LOTUS-containing MIP-1 and MIP-2 proteins. A) Live-imaging of young adults shows the distribution of LOTR-1::GFP in the presence and absence of its LOTUS and Tudor domains. B) LOTR-1 distribution during spermatogenesis in the fourth larval stage (L4). C) Live imaging of mCherry::PRG-1 and GFP::LOTR-1 in the germline L4 stage, and D) young adult stage worms. E) mCherry::PRG-1 and GFP::LOTR-1 distribution during spermatogenesis in both the presence and absence of LOTR-1’s Tudor domain. F) RNAi depletion of *mip-1* and *mip-2* in GFP::LOTR-1 compared to control RNAi. Orange boxes indicate region of spermatogenesis. Scale is 20 microns.

Tudor domains interact with PIWI Argonaute proteins via sDMAs.[76] To see if this interaction extends to *C. elegans*, the distribution of the PIWI homolog PRG-1 was examined in the presence and absence of LOTR-1, and in a strain carrying LOTR-1 with the Tudor domain deletion. Under wild-type conditions, PRG-1 is constitutively associated with germ granules from the distal germline through gametogenesis. In both *lotr-*1 mutant backgrounds, PRG-1 remains localized to germ granules during spermatogenesis, reflecting the Tudor-independent germ-granule association of LOTR-1 (Fig 2C, orange box), but is less abundant more distally (dashed arrowheads). In young adults, PRG-1 becomes expressed more distally, but its levels are reduced upon oocyte maturation in the *lotr-1* mutants (Fig 2D, dashed arrowheads). Both LOTR-1 and PRG-1 are deposited in residual bodies during spermatogenesis and absent from spermatids, in the presence or absence of the Tudor domain of LOTR-1 (Fig 2E) These findings suggest that LOTR-1 helps to recruit or stabilize PRG-1 on germ granules, but that other factors may be recruiting PRG-1 during spermatogenesis.

Next, LOTR-1 localization was examined in the absence of two other recently characterized LOTUS-containing proteins, MIP-1 and MIP-2.[71] MIP-1 and MIP-2 are <u>M</u>EG-3 interacting <u>p</u>roteins that localize to P granules throughout development and promote germ granule condensation. Although MIP-1 and MIP-2 lack Tudor domains, they each contain two eLOTUS domains that could indicate some functional redundancy with LOTR-1. RNAi depletion of *mip-1* and *mip-2* together causes the dispersal of PGL-1, PGL-3, and GLH-1.[71] Similarly, GFP-tagged LOTR-1 granules are reduced in size and less prominent around the nuclear periphery following *mip-1*; *mip-2* RNAi (Fig 2F). These results suggest that the LOTUS-domain proteins MIP-1, and MIP-2 affect the localization of LOTR-1 at germ granules.

In *Drosophila*, the LOTUS-containing protein Oskar recruits the Vasa DEAD-box helicase to germ granules, directly interacting with Vasa to stimulate its ATPase and helicase activities and promote piRNA amplification.[72,77,78] In *C. elegans*, there are two partially redundant Vasa homologs, GLH-1 and GLH-2.[79] Single mutations in either GLH are temperature-sensitive sterile, while double mutants are sterile at all temperatures. Brood size and fertility are minimally impacted in all *lotr-1* mutants at both 20°C and the first generation shifted to 26°C (Fig 3A). To test whether LOTR-1 functions synergistically with GLH-1, fertility was then examined following *glh-1* RNAi in *lotr-1* mutant backgrounds (Fig 3B). At both 20° C and 26° C, *glh-1* RNAi causes modest, yet significant (p<0.003), increases in sterility relative to wild type for all four *lotr-1* mutants tested. *glh-1* RNAi depletion reduces GLH-1::GFP expression but does not dissociate LOTR-1 from germ granules under these conditions (Fig 3C). Thus, *glh-1* and *lotr-1*, show only modest synergy in vivo, in contrast to the strong relationship between Oskar and Vasa in *Drosophila*. This difference is likely because LOTR-1 is more similar to *Drosophila* Tejas and Tapas than to Oskar, which carries a LOTUS domain but lacks the Tudor domain.

**Figure 3.**
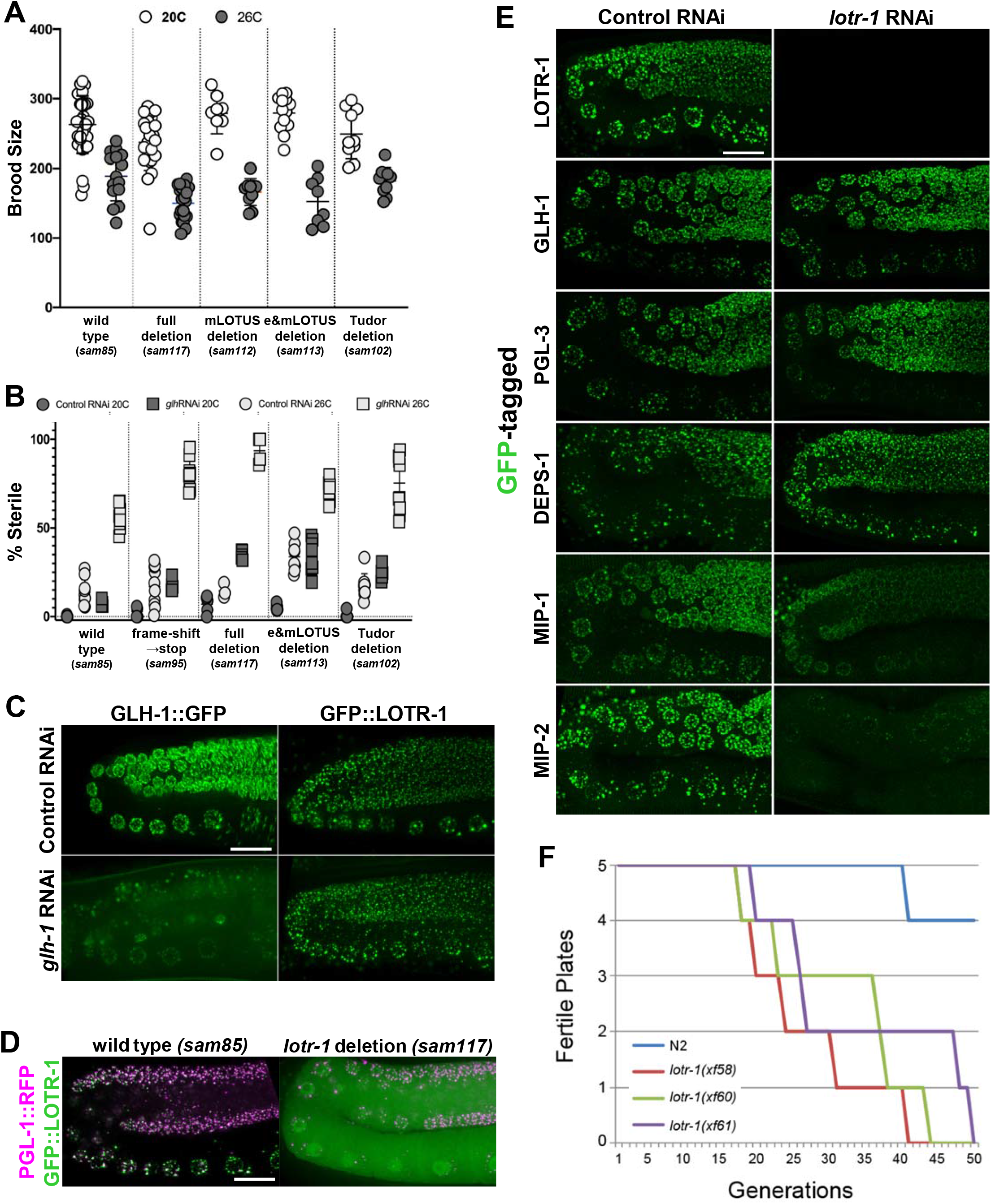
Consequences of *lotr-1* disruption. A) Fertility at permissive (20ºC) and restrictive (26ºC) temperatures in indicated *lotr-1* mutants and B) Percent sterility. C) *glh-1* RNAi in indicated *lotr-1* mutants. p<0.003. D) Live PGL-1::RFP and GFP::LOTR-1 imaging in young adult germlines. E) *lotr-1* RNAi compared to control RNAi in LOTR-1, GLH-1, PGL-3, DEPS-1, MIP-1 and MIP-2 reporter young adult worms. F) Number of fertile plates of each strain indicated per generation. The onset of sterility of *lotr-1* mutants occurred at the 18^th^ generation in the *xf58* and *xf60* alleles, and at the 20^th^ generation in the *xf61* allele. The decline in fertility proceeded in all *lotr-1* strains until no fertile plate remained. Scale is 20 microns.

We sought to address whether LOTR-1 is required for localization of other germ granule factors or vice versa. Deleting the *lotr-1* coding sequence from the 3xFLAG::GFP-tagged line (*sam117*) was not sufficient to disperse PGL-1::RFP from germ granules in distal germ cells, although PGL-1 granules are largely removed from proximal oocytes (Fig 3D). P-granule localization of GLH-1, DEPS-1, and distal PGL-3 in the adult germline is not affected by *lotr-1* RNAi (Fig 3E). However, MIP-1 and MIP-2, the two other LOTUS-containing proteins, were partially dispersed from germ granules upon *lotr-1* RNAi. This, together with the reciprocal effect of MIPs on LOTR-1 localization, suggests potential redundancy between these three LOTUS-domain containing proteins.

To explore the potential for direct associations between LOTR-1, GLH-1, MIP-1, and MIP-2, we performed yeast-two-hybrid (Y2H) analyses. MIP-1 and MIP-2 have previously been shown to both homo-and heterodimerize, as well as associate with GLH-1 through their N-terminal LOTUS domains.[71] Y2H confirmed associations between the C-terminus of LOTR-1 and both MIP-1 and MIP-2(Fig 4A, S2A Fig), whereas no interaction between LOTR-1 and GLH-1 was detected (Fig 4B, S2B Fig). The association of MIP-1 with LOTR-1 is mediated through its disordered C-terminus and the Tudor-containing C-terminal region of LOTR-1, not through their LOTUS domains. Surprisingly, Y2H results suggest that both MIP-1 and MIP-2 could interact more strongly with LOTR-1 when one or both of the LOTR-1 LOTUS domains are removed.

**Figure 4.**
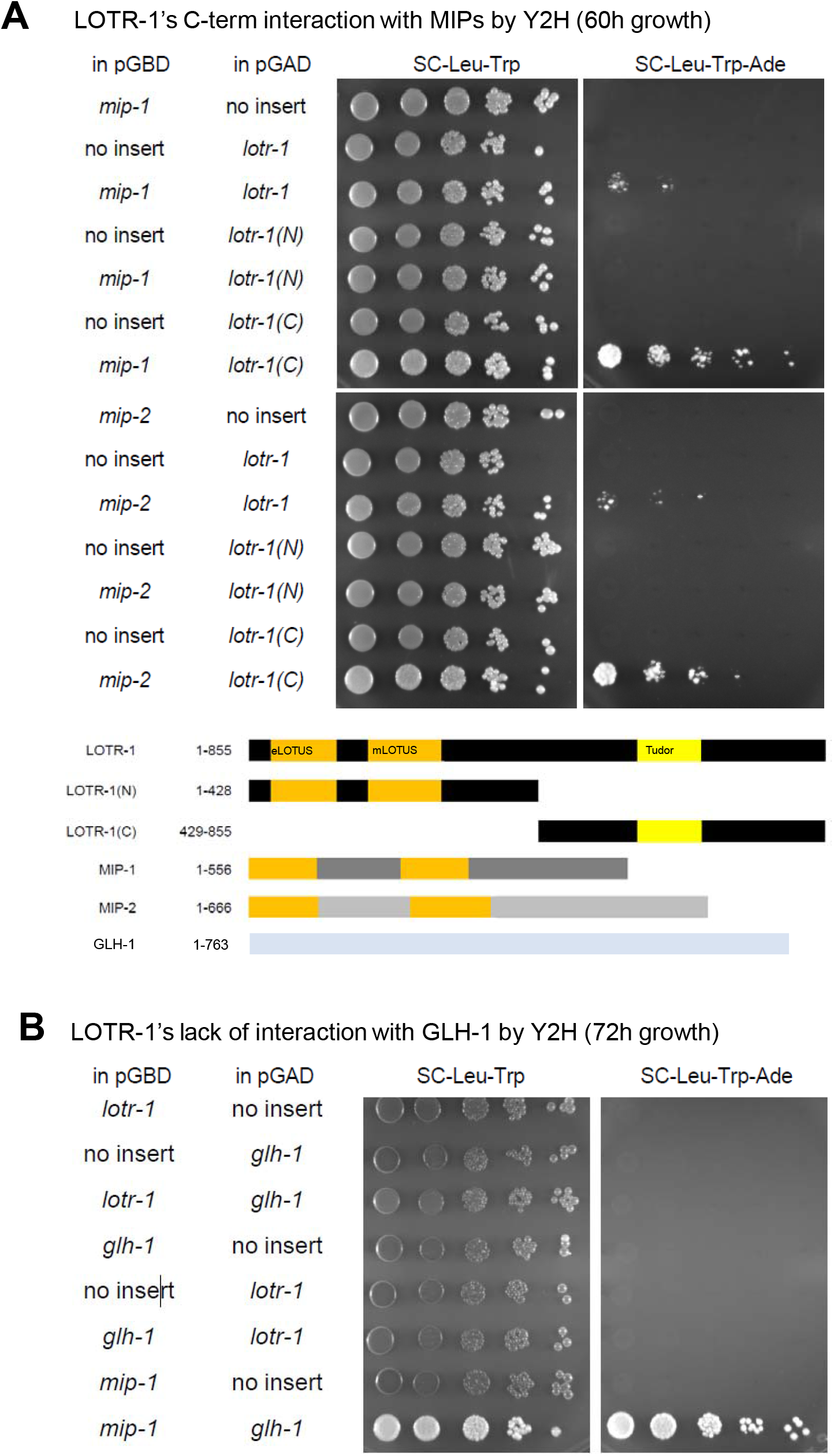
LOTR-1 interacts with MIPs but not GLH-1 by yeast-two-hybrid (Y2H). A) Association of full-length MIP-1 (top) and MIP-2 (bottom) with full-length (1-855) and C-terminal (429-855) LOTR-1 by Y2H. B) Full-length LOTR-1 and GLH-1, in both Y2H bait and prey positions, fails to demonstrate the association found between MIP-1 and GLH-1.

Mutations affecting small RNA biogenesis and amplification frequently exhibit transgenerational sterility or mortal germline (Mrt) phenotypes.[80,81] To address if LOTR-1 is required for transgenerational germline health, fertile generations were counted until sterility ensued for three different *lotr-1* mutant alleles (Fig 3F). Each of the three *lotr-1* alleles failed to propagate beyond 50 generations, while wild-type (N2) worms remained fertile. When taken together, fertility and brood size phenotypes associated with *lotr-1* are not severe and require several generations to manifest.

#### *lotr-1* mutants deregulate subsets of 22G-and 26G-RNAs

The Mrt phenotype and RNAi inheritance defects observed in *lotr-1* mutants suggest that *lotr-1* mutants are defective in some aspect of small RNA biogenesis. Since many LOTUS domain proteins have functions in piRNA biology,[76] we asked if this is the case for LOTR-1 using lines that carry a 21U-RNA/piRNA sensor construct. This transgene contains a reporter for GFP::H2B expression that has been silenced through a piRNA target site in its 3’UTR; mutations affecting piRNA biogenesis or secondary siRNA production de-silence expression of this transgene.[82] Loss-of-function alleles of *lotr-1* were crossed into two different sensor lines carrying this transgene: one in which silencing is still dependent on 21U-RNAs (S3A Fig), and another under stable RNA-induced epigenetic silencing (RNAe) that is maintained independent of the initial 21U-RNA trigger (S3B Fig). Mutations in *lotr-1* were unable to activate either piRNA sensor strain (Figs 5A-B), showing that LOTR-1 is not required for 21U-RNA-dependent silencing or for RNAe.

**Figure 5.**
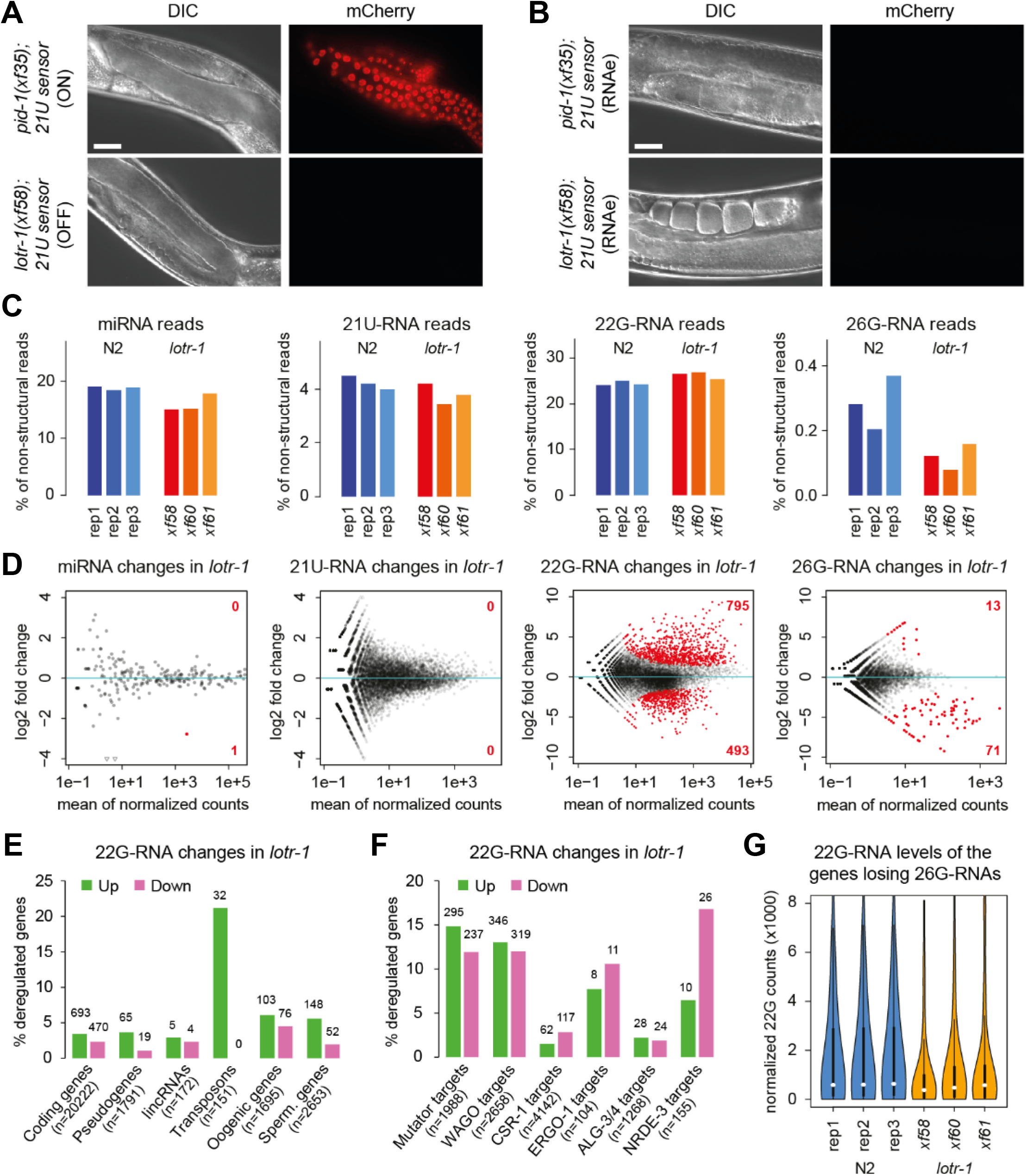
Small RNA changes in *lotr-1* mutants. A -B) Differential interference contrast, and fluorescence photomicrographs of worms carrying two different 21U sensors, one A) dependent on 21U-RNAs, and one B) under RNA-induced epigenetic silencing (RNAe). Scale is 20 microns. C) Small RNA levels for indicated populations normalized to all non-structural reads (excluding rRNA, snoRNA, snRNA, and tRNAs). D) Differential expression analysis to determine genes and transposons that are significantly depleted or enriched of mapped small RNAs in *lotr-1* mutants. The MA plots depict DESeq2 differential results for miRNAs (n=257), 21U-RNAs over 21ur loci (n=14328) and 22G-/26G-RNAs over protein coding genes (n=20222), lincRNAs (n=172), pseudogenes (n=1791) and transposons (n=151) with significant changes (>2-fold at 10% FDR) colored in red with the number of up-and down-regulated hits indicated. E-F) Bar plots depicting the number and percentage of genes with significantly deregulated 22G-RNAs in the indicated gene classes and target lists from previous studies. G) Violin plot showing the distribution of normalized 22G-RNA levels of the 71 genes with significant 26G-RNA depletion (p=0.015 using a two-sided Wilcoxon signed-rank test for a group-wise comparison).

As described in the introduction, *C. elegans* expresses a variety of small RNA species.[80] To address if LOTR-1 is required for some aspect of small RNA biogenesis, we sequenced small RNAs in wild-type and *lotr-1* mutants. Small RNA sequencing in gravid adults showed that 21U-RNAs and miRNAs are hardly affected in *lotr-1* mutants, while 26G-RNAs are partially depleted (Fig 5C, D). Global 22G-RNA levels are not changed overall; however, hundreds of genes have deregulated 22G-RNA levels in *lotr-1* mutants (Fig 5C-D, S3 Table). Next, we asked if the deregulated 22G-RNAs map to particular gene classes. In *lotr-1* mutants, 22G-RNAs are deregulated in similar proportions in coding genes, pseudogenes, lincRNAs, oogenic and spermatogenic genes (Fig 5E). While no transposons show depletion of 22G-RNAs, 21% of them display upregulated 22G-RNA levels in *lotr-1* mutants. Aggregated data for Tc1 transposons show 3-fold increased 22G-RNA levels in *lotr-1* mutants, but with a marginal FDR-adjusted p-value of 0.15 due to replicate variation (S3 Table). A reporter for Tc1 transposon mobility was not affected by the *lotr-1* mutation, suggesting that the upregulation of Tc1 22G-RNAs observed in *lotr-1* animals does not disrupt Tc1 silencing (S3C-E Figs). After separating 22G-RNAs into subgroups according to their Argonaute cofactors and target genes [80], we find that most genes with increased 22G-RNA levels in *lotr-1* mutants belong to the mutator and WAGO classes (Fig 5F). Conversely, a similar proportion of mutator, WAGO, ERGO-1, and NRDE-3 targets are depleted of 22G-RNAs in *lotr-1* mutants. Overall, these results suggest that hundreds of genes show unbalanced 22G-RNA expression in *lotr-1* mutant animals across a range of gene classes and 22G-RNA subgroups.

Seventy-one genes seem to be depleted of 26G-RNAs (Fig 5D). Of these, all but nine are annotated to be targets of the known effectors ERGO-1, NRDE-3, or ALG-3/4 (S3 Table).[83–85] The depletion of 26G-RNAs in these 71 genes in *lotr-1* mutants is accompanied by a tendency to deplete downstream 22G-RNAs (Fig 5G). A GFP::NRDE-3 transgene, which localizes from the nucleus to the cytoplasm upon disruption of the 26G-RNA pathway,[84,86] remained expressed in the nucleus when crossed into a *lotr-1* mutant (S3F-G Fig), indicating the 26G-RNA pathway is not critically impaired in *lotr-1* mutants. Taken together, these results suggest that while LOTR-1 is not absolutely required for normal 26G-RNA biogenesis and silencing, a subset of 26G-RNA targets is affected by the depletion of LOTR-1.

### LOTR-1 binding partners include components of germ granules and the cytoskeleton

We sought to understand how LOTR-1 regulates small RNA biogenesis by finding LOTR-1 binding partners. The N-terminal 3xFLAG tag introduced into the endogenous LOTR-1 locus was used to immunoprecipitate LOTR-1 from both young adults and embryos for quantitative mass spectrometry (IP-qMS). To determine differential enrichment over a control, CRISPR-Cas9 genome editing was used to delete the *lotr-1* coding sequence so that the endogenous *lotr-1* promoter and 3’UTR drive expression of 3xFLAG::GFP, and anti-FLAG IP-qMS from the *lotr-1* deletion was compared to wild-type LOTR-1 (Fig 6A). LOTR-1 IP-qMS experiments were performed in quadruplicate for each IP sample on two different occasions (Round 1 and Round 2) (Fig 6B). Including LOTR-1, eight proteins were enriched in LOTR-1 IPs in all datasets, both in embryos and young adults (Figs 6A-C). These include the germ granule-associated Argonaute proteins PRG-1 and WAGO-1, the helicase ZNFX-1, the 3’UTR cleavage and stimulation factors SUF-1 and CPF-1, the histone deacetylase SIR-2.2, and F46G10.1. These eight proteins represent a central core of LOTR-1 interactions, reinforcing the role of LOTR-1 in germ granules and small RNA biogenesis.

**Figure 6.**
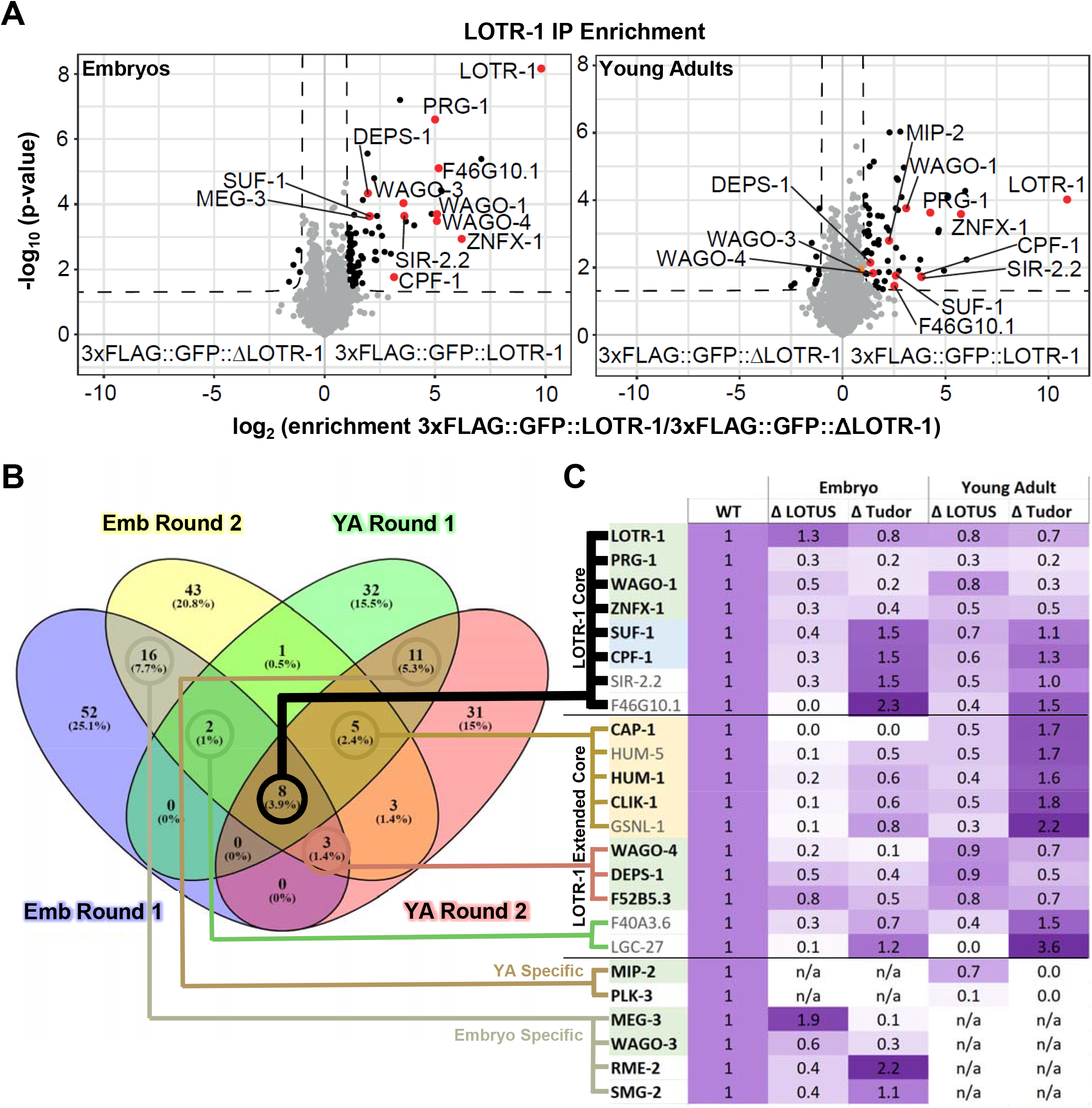
LOTR-1 protein associations. A) Volcano plots show the significance and enrichment of proteins that immunoprecipitated with 3xFLAG::GFP::LOTR-1 over the *lotr-1* deletion expressing GFP::3xFLAG alone, as identified by qMS. B) Venn diagram shows significantly enriched proteins that overlapped between two rounds of LOTR-1 IP-qMS from both embryo and young adult lysates. C) Heat map showing the change in LOTR-1 association in embryos and young adults when LOTUS or Tudor domains of LOTR-1 were deleted.

Expanding the interaction list by an additional ten proteins that showed significant enrichment in at least three out of the four embryo and young adult IP-qMS experiments, now includes the germ-granule proteins DEPS-1, the WAGO-4 Argonaute, and the YHTDC2-like DExH-helicase (F52B5.3) that was previously shown to interact with the germ-granule Argonaute CSR-1.[87] Of these 18 LOTR-1-interacting proteins, 5 are known to bind actin and regulate the cytoskeleton, including CAP-1, HUM-1, HUM-5, CLIK-1, and GSNL-1.[88–91] 12 of the 18 proteins originate from transcripts abundantly expressed in the germline (Fig 6C, bold), while transcripts encoding the remaining six (HUM-5, GSNL-1, F40A3.6, SIR-2.2, F46G10.1, and LGC-27) are lowly expressed in the germline and may reflect LOTR-1 interactions in whole-worm or embryo lysates that are not replicated *in vivo*.[92,93]

To distinguish if LOTR-1 associations obtained by co-immunoprecipitation (co-IP) are facilitated through its LOTUS or Tudor domains, LOTR-1 carrying a deletion of either the e+mLOTUS (*sam113*) or Tudor (*sam102*) domains was immunoprecipitated from embryos and young adults and then analyzed with qMS. Deleting the LOTUS domains had a more pronounced effect on the association of LOTR-1 with cytoskeletal proteins, 3 ‘UTR-cleavage and stimulation factors, and four proteins that were not readily grouped (Fig 6C, orange, blue, white). Associations with other germ-granule proteins were impacted by both LOTUS and Tudor deletions (Fig 6C, green), and the effect was slightly more pronounced in the absence of the Tudor domain, which results in the dispersal of LOTR-1 from germ granules.

Of the LOTR-1 associations specific to both embryo IPs, MEG-3 and WAGO-3 stand out because of their previously described association with germ granules. The association of LOTR-1 with MEG-3 in embryos is dependent on the LOTR-1 Tudor domain, and it is enhanced in the absence of its LOTUS domains (Fig 6C, S1 Table). This may suggest that Tudor-dependent interactions between MEG-3 and LOTR-1 during embryogenesis are kept in check by other associations mediated through its LOTUS domains. In turn, other LOTR-1 interactions are more prominent in young adults, including, for example, those with cytoskeletal proteins (Fig 6C, S1 Table). The LOTUS-domain containing MIP-2 protein associates with LOTR-1 in the young adult germline and this interaction is dependent on the Tudor domain, consistent with our Y2H results (Fig 4A). Another of the more pronounced LOTR-1 interactors specific to young adults is the polo-like kinase PLK-3, and this interaction is disrupted when either the LOTUS or Tudor domains of LOTR-1 are removed. The significance of these embryo and young adult-specific interactions warrants additional attention.

### LOTR-1 functions with ZNFX-1 to distribute WAGO and mutator-class 22G-RNAs along germline RNAs

As proteins bearing LOTUS and Tudor domains, like Oskar, interact with helicases, we reasoned that the strong association identified by IP-qMS with the ZNFX-1 helicase may illuminate LOTR-1 function *in vivo*. In the adult germline, ZNFX-1 is positioned adjacent to P granules and defines a sub-compartment of germ granules called Z granules.[69,70] Unlike the offset/docking observed between LOTR-1 and PGL-1 (Figs 1D-E), we found that RFP::ZNFX-1 and GFP::LOTR-1 are not offset in adult germ cells, but overlap (r=0.82 ±0.03), suggesting that LOTR-1 is also a component of Z granules (Fig 7A). An interaction between ZNFX-1 and LOTR-1 was also supported by Y2H (S2C Fig). Deleting the *lotr-1* coding region causes most ZNFX-1 to disperse from germ granules, primarily in oocytes in the proximal adult germline (Fig 7B, dashed arrows). Proper ZNFX-1 localization is not dependent upon the LOTR-1 LOTUS domains but rather on its Tudor domain (Fig 7B). An early frameshift mutation was introduced into ZNFX-1 to determine whether loss of ZNFX-1 impacts GFP::LOTR-1 and RFP::PRG-1 expression. In the first generation of homozygous *znfx-1* animals, LOTR-1 and PRG-1 were localized in germ granules as previously observed (Fig 7C). This could be due to maternal rescue, as LOTR-1 expression is more diffuse in the second *znfx-1* generation toward the most distal and proximal regions of the germline. PRG-1 was mostly diffuse and unattached from germ granules (dashed arrows) in second-generation *znfx-1* mutants, but also consistently showed some formation of larger aggregates. In *lotr-1; znfx-1* double mutants, PRG-1 dispersal was generally impacted to the same degree as *lotr-1* and *znfx-1* single mutants (Fig 7D). These results show that ZNFX-1 and LOTR-1 reciprocally affect each others’ localization, suggesting that these two Z-granule proteins act in the same pathway.

**Figure 7.**
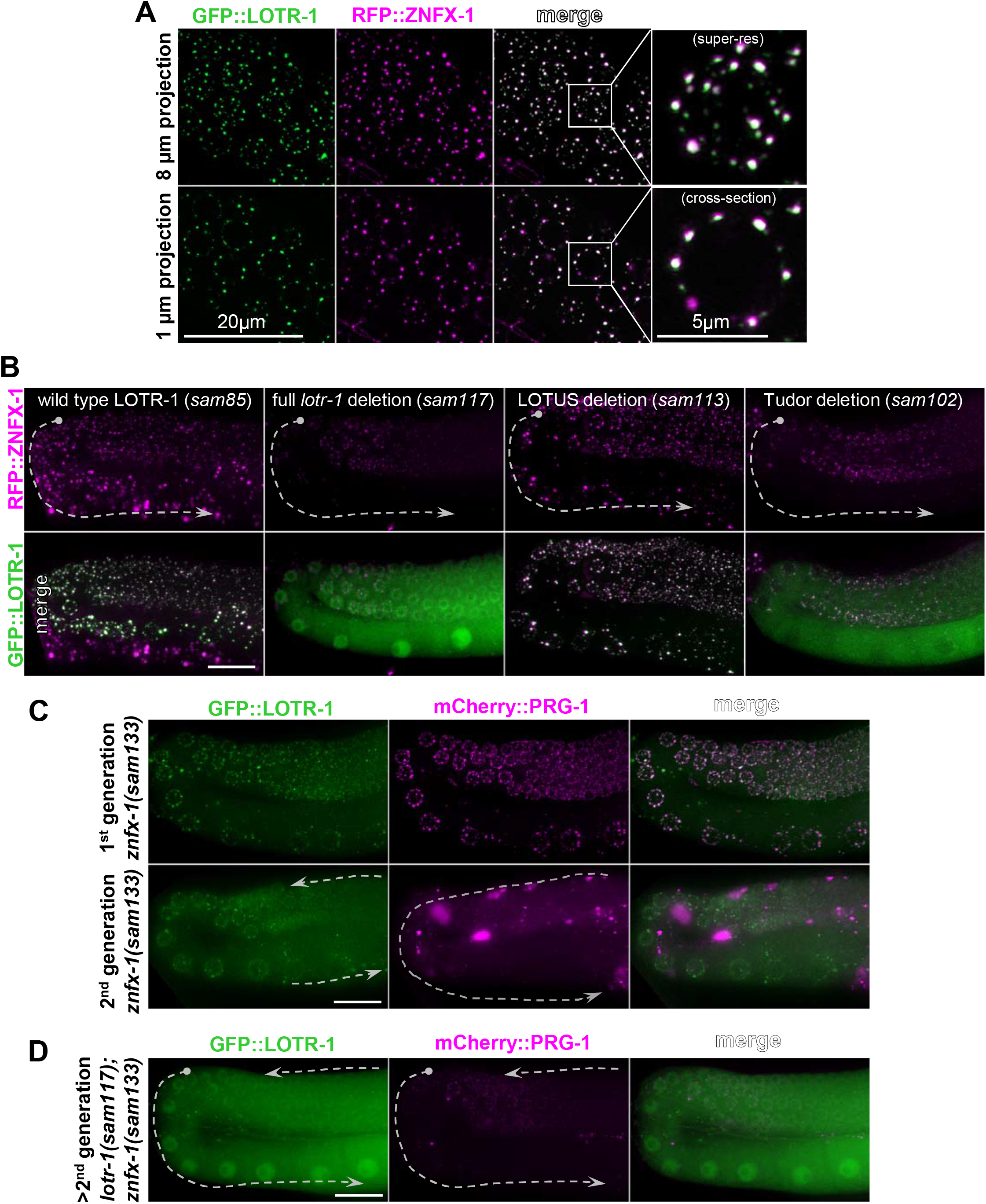
LOTR-1 localizes to and impact Z granule composition. A) Live super-resolution confocal imaging of RFP::ZNFX-1 and GFP::LOTR-1 in pachytene cells of the adult germline. B) Live imaging of RFP::ZNFX-1 and GFP::LOTR-1 in the germline of wild-type and *lotr-1* mutant adults. C) Comparison of GFP::LOTR-1 and mCherry::PRG-1 expression in the germlines of first and second generation of *znfx*-mutant worms. D) mCherry::PRG-1 expression in *lotr-1; znfx-1* double mutants. Scale is 20 microns unless otherwise stated.

ZNFX-1 has been suggested to promote balanced production of 22G-RNAs across target transcripts by positioning the RdRP EGO-1 toward the 3’ end of RNAi target transcripts.[69,70] In *znfx-1* mutants, 22G-RNA coverage is altered, revealing a shift toward the 5’ end of the target mRNAs.[70] Metagene analysis of all WAGO and mutator targets showed a similar 5’ shift of 22G-RNAs in three different *lotr-1* mutant alleles compared to wild-type replicates (Fig 8A-B). Unlike *znfx-1*, the 5’ shift in 22G-RNA coverage in *lotr-1* mutants is observed in WAGO/mutator, but not in CSR-1 targets (Fig 8A-C). This trend is more pronounced in mutator targets with upregulated 22G-RNA levels (S4 Fig A-B). 22G-RNA coverage of ALG-3/4 and ERGO-1 targets was not affected in *lotr-1* mutants (S4 Fig C-D). Notably, CSR-1 was not enriched in LOTR-1 IP-qMS, in either the embryo or young adult lysates. Therefore, we conclude that LOTR-1 functions with ZNFX-1 to balance the production of 22G-RNAs across WAGO/mutator, but not CSR-1, target RNAs.

**Figure 8.**
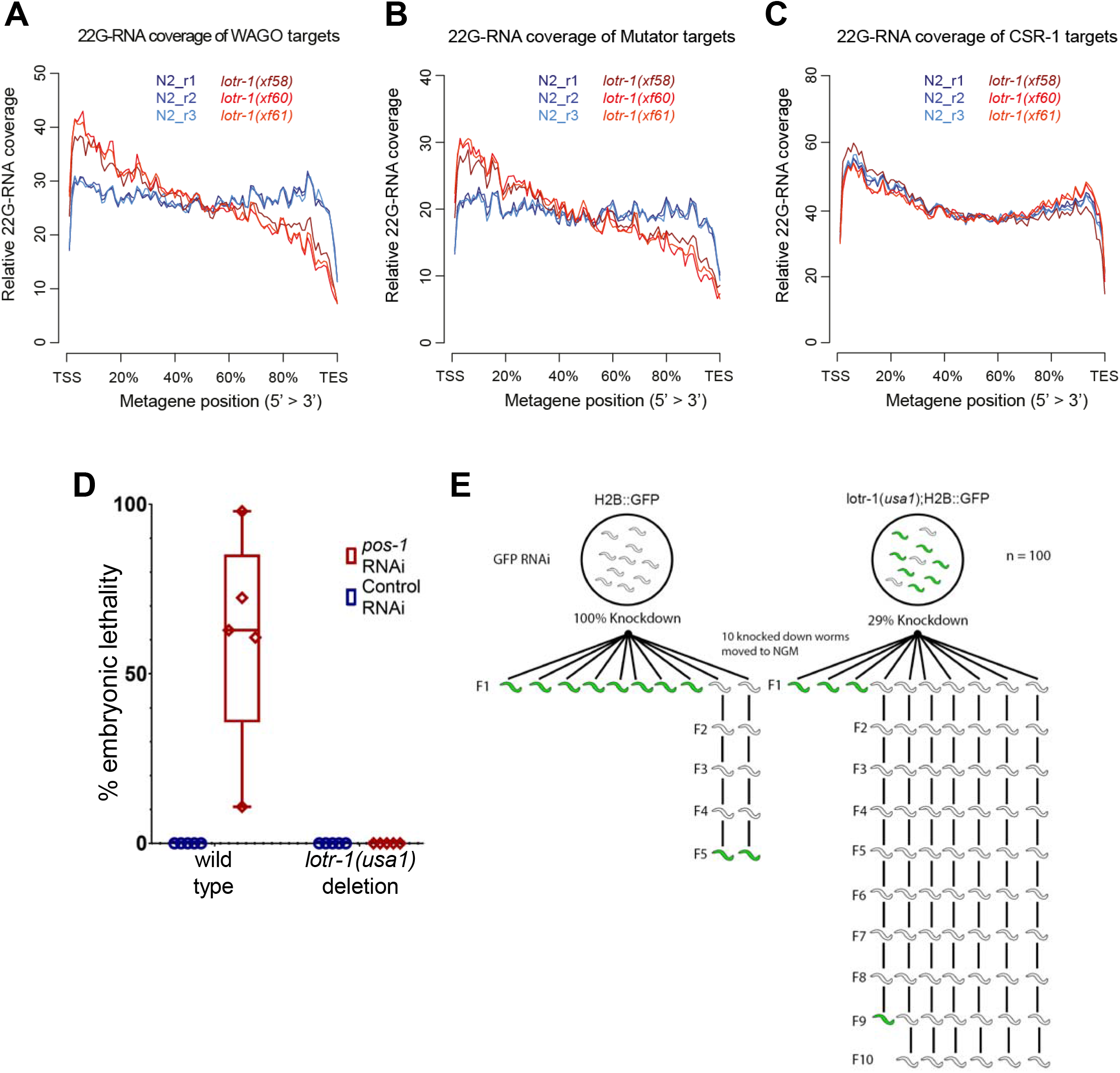
Altered 22G-RNA distribution and RNAi defects in *lotr-1* mutants. Metagene plots to visualize relative 22G-RNA distribution in wild-type (N2) and *lotr-1* mutants over A) WAGO targets, B) mutator targets, and C) CSR-1 targets. D) Resistance to *pos-1* RNAi-induced embryonic lethality in a *lotr-1* mutant. E) RNAi inheritance in a *lotr-1* mutant. Knock-down of GFP persists over more generations in the *lotr-1* deletion.

The imbalance of 22G-RNA coverage over WAGO/mutator targets observed in *lotr-1* mutants, and the interaction of LOTR-1 with ZNFX-1 and WAGO proteins, may point to defects in exogenous RNAi and its inheritance. To test whether this is the case, the incidence of embryonic lethality following *pos-1* RNAi was examined in both wild type and *lotr-1* mutants (Fig 8D). Embryonic lethality was completely suppressed in all five *lotr-1* replicates. Additionally, to test for defects in the inheritance of RNAi, GFP RNAi was performed in wild type and *lotr-1* mutants carrying an H2B::GFP reporter (Fig 8E). GFP RNAi was 100% effective in knocking down GFP expression in wild-type worms, but only knocked down expression in 29% of *lotr-1* mutants, in line with the resistance to *pos-1* RNAi (Fig 8D). To determine how quickly GFP expression was restored in silenced wild-type and *lotr-1* worms, ten knocked down worms from each strain were transferred from RNAi to non-RNAi plates and examined for expression in the following generations. While GFP expression was restored within one to five generations in wild-type worms, most *lotr-1* mutants maintained GFP silencing past ten generations (Fig 8E). Altogether these findings suggest that exogenous RNAi and its inheritance across generations are compromised in *lotr-1* mutants, which may be related to the imbalance in the distribution of 22G-RNAs across target transcripts.

To confirm whether ZNFX-1 interactions reflect the core set of proteins bound to LOTR-1, anti-FLAG IP-qMS was performed on embryos and young adults expressing 3xFLAG::GFP::ZNFX-1 or an untagged control (Fig 9A). In addition to ZNFX-1, proteins enriched over untagged control in both embryo and young adult included LOTR-1 and its interactors (i.e., PRG-1, DEPS-1, WAGO-1, WAGO-4 and RME-2). SMG-2, WAGO-3, and the 3’ UTR cleavage and polyadenylation factors SUF-1 and CPF-1 were enriched in embryos only, while MIP-2 and PLK-3 were only enriched in young adults. These results support a substantial overlap of interactions between ZNFX-1 and LOTR-1 within Z granules. In *lotr-1(xf58)* mutant embryos, the association of ZNFX-1 with PRG-1, DEPS-1, WAGO-1, WAGO-3 and WAGO-4 is reduced by about 50%, while its association with SUF-1, CPF-1, and SMG-2 is practically eliminated (Fig 9B-C, S2 Table). This suggests that LOTR-1 stabilizes ZNFX-1 interactions with Argonaute proteins while acting as the link between ZNFX-1 and mRNA/3’UTR binding factors. In *lotr-1* mutant adults, the association of ZNFX-1 with PLK-3 is reduced 80%, suggesting that LOTR-1 is the link here as well. Interestingly, the LOTUS-containing protein MIP-2 increases its association with ZNFX-1 in the *lotr-1* mutant, which may support a compensatory function for MIP-2 and LOTR-1.

**Figure 9.**
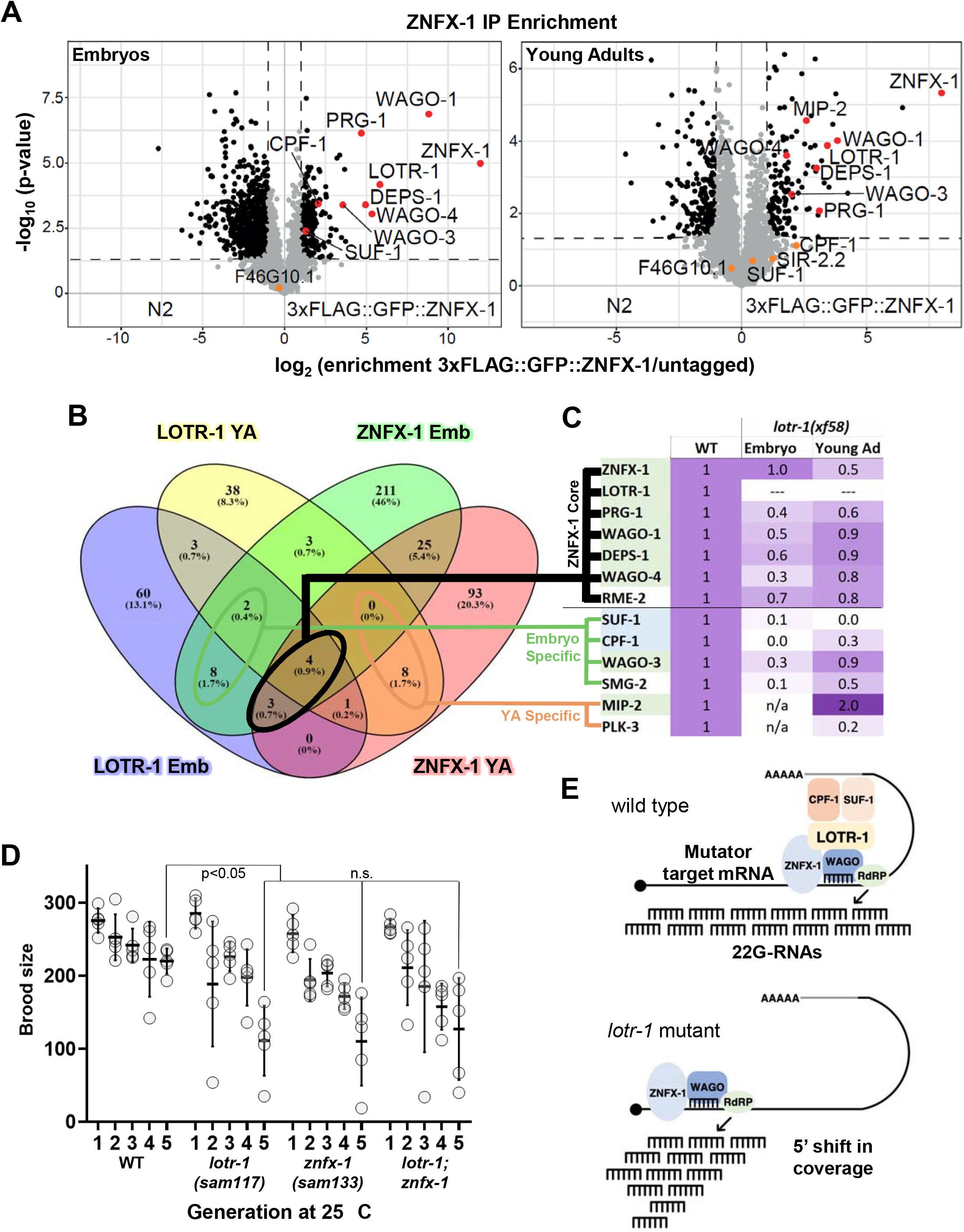
ZNFX-1 protein associations. A) Volcano plots show the significance and enrichment of proteins immunoprecipitated with 3xFLAG::GFP::ZNFX-1 over untagged ZNFX-1, as identified by qMS. B) Venn diagram shows significantly enriched proteins that overlapped between LOTR-1 and ZNFX-1 IP-qMS from both embryo and young adult lysates. C) Heat map showing the change in ZNFX-1 association when *lotr-1* was mutated. D) Brood sizes at 25ºC in *lotr-1* and *znfx-1* mutants over five generations. A T-test was used to determine assess significance at generation five. E) Model of LOTR-1 function in Z granules. LOTR-1 could tether 3’ end processing factors CPF-1 and SUF-1 to RdRP-dependent amplification of 22G-RNAs, keeping 22G-RNA distribution balanced. In the absence of LOTR-1, RdRP activity shifts to the 5’ end of WAGO/mutator targets.

ZNFX-1 and LOTR-1 colocalize within Z granules (Fig 7A), balance production of 22G-RNAs across their targets (Fig 8), have overlapping binding partners (Figs 6 and 9), and *lotr-1; znfx-1* double mutants have a similar impact on PRG-1 dispersal as the single mutants (Figs 2D and 7C-D). However, some subtle differences in ZNFX-1 and LOTR-1 association point to distinct roles. For example, LOTR-1 has a stronger affinity for cytoskeletal proteins while ZNFX-1 has a stronger affinity for a handful of mitochondrial oxidoreductase proteins, and in *lotr-1* mutants the 5’ shift in 22G-RNA coverage is specific to WAGO and mutator targets. Because *znfx-1* and *prg-1* mutants have transgenerational epigenetic inheritance defects that manifest after several generations at 25° C [69,94], brood sizes of single and double *lotr-1; znfx-1* mutants were compared to wild-type broods over the course of five generations (Fig 9D). Again, no additive effect was observed between the double and single mutants. These results add support to the hypothesis that LOTR-1 and ZNFX-1 are integral to one another’s primary function in Z granules – both acting to balance 22G-RNA distribution across WAGO/mutator targets (Fig 9E).

## Discussion

Considering the central role of *Drosophila* Oskar and other LOTUS domain-containing proteins like Tejas/TDRD5 and Tapas/TDRD7 in germline specification and development, we sought to ask whether LOTUS domain proteins in *C. elegans* function similarly and could thus be used to model TDRD5 and TDRD7 function in mammals. We have shown that LOTR-1 is the only protein in *C. elegans* where extended and minimal LOTUS domains are paired with a C-terminal Tudor domain, suggesting homology with TDRD5 and TDRD7.

We found that LOTR-1 colocalizes and coimmunoprecipitates with the Z-granule protein ZNFX-1 and that the two proteins interact in yeast two hybrid (Y2H) assays (Figs 6-7, 9 and S2C). Associations between the two proteins are maintained through LOTR-1’s Tudor domain, while LOTUS domain mutations do not impact the Z-granule localization of LOTR-1. *lotr-1* mutants have reduced levels of 26G-RNAs (Fig 5) and, while global 22G-RNA levels remain the same, these are deregulated and altered within specific subpopulations. Specifically, 22G-RNA levels decrease on some genes that lose 26G-RNAs in *lotr-1* mutants and, like *znfx-1* mutants, 22G-RNA coverage of WAGO/mutator targets show a 5’ shift, particularly on upregulated transcripts (Figs 5, 8). However, unlike *znfx-1*, in *lotr-1* mutants this 5’ shift extends only to WAGO/mutator targets and not to CSR-1 targets. This work suggests that LOTR-1 helps balance 22G-RNA distribution across WAGO/mutator targets to preserve germline integrity and fertility across generations.

One outstanding question is the functional relationship between LOTR-1, which combines its two LOTUS domains with a Tudor domain, and the recently discovered paralogs MIP-1 and MIP-2, which each contain two LOTUS domains and long intrinsically disordered regions (IDRs). Y2H assays show that both MIP-1 and MIP-2 associate with GLH-1/Vasa[71], while Y2H does not support a direct interaction between LOTR-1 and GLH-1 (Figs 4B and S2B). The association of MIP-1 and MIP-2 with GLH-1 in *C. elegans* is likely similar to Oskar’s association with Vasa in *Drosophila*.[72,77] Interestingly, we show that the C-terminus of LOTR-1 also associates with the C-terminus of both MIP-1 and MIP-2 via Y2H (S2 Fig), and that MIP-2 is enriched in both ZNFX-1 and LOTR-1 IP-qMS from young adult lysates (Figs 6, 9). Depletion of LOTR-1 or either of the MIPs can reciprocally impact each other’s association with germ granules (Figs 2E and 3E). This complicates labeling these proteins as strict components of either P-or Z-granules, implying a degree of inter-granule crosstalk. It also suggests there may be partial redundancy among these three LOTUS-domain proteins. For example, MIP-2 increases its association with ZNFX-1 in *lotr-1* young adults, implying that it can partially compensate for the absence of LOTR-1. It remains to be determined whether the association of MIP-1 and MIP-2 with GLH helicases in P granules is analogous to the association of LOTR-1 with the ZNFX-1 helicase in Z granules, and whether these P-and Z-granule associations distinguish 22G-RNA distribution across CSR-1 and WAGO/mutator targets.

Loss of LOTR-1, either by deletion or RNAi depletion, failed to impact GLH-1 and DEPS-1 distribution but reduced PRG-1, PGL-1, and PGL-3 foci in proximal oocytes (Figs 2D, 3C-E). GLH-1, DEPS-1, PRG-1, PGL-1, and PGL-3 are all constitutive P-granule components, and thus far they have not been observed to colocalize with Z granules; however, both PRG-1 and DEPS-1 were significantly enriched by LOTR-1 and ZNFX-1 IP-qMS (Figs 6, 9). While this association may reflect associations in early embryo germline blastomeres, before P-and Z-granule demixing in primordial germ cells (PGCs) [69], it strengthens the likelihood of dynamic inter-granule crosstalk in PGCs and in germ cells during larval and adult development. Interestingly, WAGO-3 and SMG-2 associations with both ZNFX-1 and LOTR-1 are only significant in embryo lysates, despite their abundant mRNA expression in adults (Figs 6, 9). In contrast, the association between LOTR-1 and MEG-3 corresponds with MEG-3 expression in early embryos. MEG-3 is a scaffold protein that nucleates germ-granule assembly after fertilization in the one-cell zygote.[95–98] The interaction with MEG-3 is dependent on the Tudor domain of LOTR-1 and implicates MEG-3 in the initial localization of LOTR-1 to germ granules. The LOTUS domains of LOTR-1 may keep the MEG-3 association in check, as the LOTR-1 association with MEG-3 increases when the LOTUS domain is deleted.

Additional insight into the contribution of LOTR-1’s LOTUS and Tudor domains can be inferred from domain-specific deletions. Co-IP data show that the association of germ granule proteins PRG-1, WAGO-1, WAGO-4, DEPS-1 and ZNFX-1 with LOTR-1 in vivo is lost upon deletion of either its LOTUS or Tudor domains. The latter result is consistent with the observation that LOTR-1 is dispersed from germ granules in the absence of the Tudor domain (and thus less likely to encounter other germ granule components); however, LOTR-1 localization to P granules is independent of its LOTUS domains. This suggests that the association of LOTR-1 with these P-granule proteins depends at least partially on specific interactions with its LOTUS domains. LOTR-1 associations that depend specifically on its LOTUS domain, based on co-IP data, include the actin-binding proteins HUM-1, HUM-5, CLIK-1, and GSNL-1, as well as 3’UTR associated proteins SUF-1 and CPF-1 (Fig 6C). Therefore, the LOTUS domain may be used to tether LOTR-1 to the cytoskeleton and 3’UTR of Z granule-associated transcripts, providing potential mechanisms for Z-granule demixing or the ability to counter the 5’ coverage bias of WAGO 22G-RNA targets during the amplification of small RNAs. In fact, the most significant impact on Z-granule composition in a *lotr-1* mutant is decreased association of SUF-1 and CPF-1 (Fig 9A, C). This suggests that RdRP-dependent 22G-RNA amplification along WAGO/mutator targets may be enriched within the more 3’ regions of the targeted transcript by the interaction between LOTR-1 and the 3’ UTR.

In *Drosophila, tejas/tapas* double mutant males are infertile.[99,100] Similarly, in mice, loss of either TDRD5 or TDRD7 will cause male-specific sterility, with defects during spermiogenesis.[31,32,101,102] Even though LOTR-1 does not affect piRNA biogenesis in *C. elegans*, our study reveals an intriguing parallel to mouse mutants lacking TDRD5. The loss of 22G-RNAs from the 3’ regions of 22G-RNA producing loci in *lotr-1* mutants resembles the loss of piRNAs from the 3’ regions of (pachytene) piRNA producing loci in Tdrd5 mutant mice.[98] It will be interesting to test if mammalian ZNFX1 acts together with TDRD5, even to the extent of producing Z-granule-like molecular condensates, and to resolve the mechanism that maintains small RNA production over the complete length of small RNA producing loci. Given our identification of 3’ processing factors in association with LOTR-1 and ZNFX-1, an intriguing hypothesis would be that LOTR-1 and ZNFX-1 target the small RNA producing machinery, be it RdRP in *C. elegans*, or specific nucleases in mammals, to the 3’ ends of transcripts through interactions with 3’ end processing complexes (Fig 9D).

## Materials and Methods

### Sequence and Structural Analysis

To characterize LOTR-1 and its interactions, we used sequence analysis (PSI/Delta-BLAST, HHPred, and multiple sequence alignment with PRALINE) to identify conserved protein domains.[103–106] Homology modeling was performed with RaptorX: eLOTUS domain was constructed from Oskar LOTUS (PDB ID: 5NT7); mLOTUS domain from Cdt1 (PDB ID: 5MEC), which contains winged-helix domains that interact with subunits of the MCM helicase motor; and Tudor domain from TD3 (PDB ID: 4b9wA) from TDRD1.

### Strain Generation & Maintenance

*C. elegans* strains were maintained using standard protocols.[107] A complete list of strains and alleles generated for this study is provided (S4 Table). CRISPR/Cas9 was used to place tags on endogenous genes and generate mutant alleles as described.[108] All alleles generated for this study were sequence verified.

### Imaging

Live worms were mounted on agarose pads in egg buffer (25 mM HEPES, 120 mM NaCl, 2 mM MgCl2, 2 mM CaCl2, 50 mM KCl, and 2 mM levamisole) and imaged with a 63X/1.40 oil objectives under fixed exposure conditions. Figs 1D, 2, 3C-E, and 7B-D are deconvolved 30μM projections of the germline loop region acquired using Leica AF6000LX acquisition software on an inverted Leica DMI6000B microscope with an attached Leica DFC365FX camera. Figures 1E and 7A are 8μM and 1μM projections of pachytene germ cells acquired using Zen Blue 3.1 acquisition software on an upright Zeiss LSM 980 with AiryScan2 processing to achieve up to 120nm super-resolution.

### RNAi

RNAi feeding was performed as previously described.[109] The L4440 plasmid in HT115 bacteria was used as the RNAi control. *glh-1* and *pgl-1* RNAi were performed starting on L4 worms, with at least three biological replicates, and used feeding constructs previously described.[110] To deplete *mip-1* and *mip-2*, equal parts from the Ahringer RNAi feeding library clones were mixed and fed to L4 worms.

RNAi resistance experiments were performed by placing five L1 larvae from wild-type N2 and *lotr-1(usa1)* mutants on *pos-1*(RNAi) or control L4440(RNAi) plates at 20°C. After 24 hours post-L4 stage, adults were removed, and eggs were allowed to hatch for 24 hours before scoring larvae and unhatched eggs. For GFP knockdown and transgenerational RNAi inheritance, L1 animals expressing H2B::GFP in wild-type N2 and *lotr-1(usa1)* mutants were placed on GFP(RNAi) and control L4440(RNAi) plates at 20°C (in triplicate) and 100 progeny from each plate were scored for GFP expression. From the GFP(RNAi) plates, ten silenced progeny from both wild type and *lotr-1* mutants were singled to individual plates seeded with OP50 (referred to as “P0” generation in NGM). For each line, all progeny (n>50) either showed silencing or were unsilenced. Unsilenced lines were propagated and examined for GFP expression for a maximum of ten generations.

### Brood Size

Brood sizes were counted from each strain for Figure 3A after placing ten or more L4 worms on individual plates at 20°C and 26°C. For the generational brood size assay in Figure 9D, 5 L4 worms from each strain were cloned to individual plates and shifted from 20°C to 25°C. Worms were kept at 25°C for 5 generations, with L4 progeny from each plate used to start the subsequent generation.

### Fertility

For each strain the fertility was determined by plating L4 worms at both 20° and 26°. Hatched F1 progeny were then picked to 10 plates with 25 worms on each plate. The percent of grotty (uterus filled with unfertilized oocytes and terminal embryos) and clean (germline atrophy with empty uterus) sterile F1s were scored when they reached day 2 of adulthood.

### Germline Mortality

Worms were assessed for the mortal germline phenotype using the assay described in Ahmed et al. 2000.[111] Three *lotr-1* strains (with the *xf58, xf60, xf61* alleles) and wild-type N2 animals were synchronized by bleaching and overnight hatching in M9 and L1-arrested animals. Five L1-L2 worms were picked to a new plate and grown at 25°C. The sixth day L1-L2 worms, corresponding to the third generation from the picked worms, were again hand-picked to a new plate. Fertile generations were counted until sterility ensued and no progeny could be isolated. Five replicates were used in this assay, N2 worms were used as control. Worms were always picked before starvation to avoid any effect it might cause during the assay, including extension of transgenerational germline lifespan.[112]

### Yeast Two Hybrid

Full-length or truncated cDNAs of *mip-1, mip-2, glh-1* and *lotr-1* and DNA encoding 3xFLAG tag were cloned in-frame with *GAL4-AD* or *GAL4-BD* into the multiple cloning site of pGAD-C1 and pGBD-C1.[113] Yeast cells from strain PJ69-4a were transformed with 1ug of plasmid and carrier DNA with lithium acetate.[114] Transformed cells were then plated on the appropriate drop-out media, and the presence of the newly introduced plasmids was checked by colony PCR. Bait-prey interactions were done on SC-Leu-Trp-Ade plates or SC-Leu-Trp-His plates supplemented with varying concentrations of 3-amino-1,2,4-triazole, as previously described.[71]

### Reporters for small RNA activity

Experiments were performed as previously described.[82,84,86,115] Crosses (S3 Fig) were set up in 6 cm Petri dishes with only 10 μl of OP50 to favor nematode meeting and mating. Wide-field fluorescence microscopy images (Figs 5A-B, S3F-G) were acquired and processed with the Leica Application Suite (LAS) software (v3.1.0) on a Leica DM6000B.

### Tc1 Reversion

Tc1 reversion experiments were performed as previously described.[116] Crosses were set up, and cross-progeny genotyped as described for the sensor experiments above. Twitcher *unc-22(st136)* worms homozygous for *lotr-1* or for *wago-1/2/3* mutations were grown at 25°C. 24 independent populations of *lotr-1; unc-22(st136)* per *lotr-1* allele. Both controls and *lotr-1* populations were grown in parallel for several generations, namely four plate passages, corresponding to 8-12 generations. The N2, and *wago-1/2/3* mutant controls were grown in 2 independent populations for the same duration of time. Plates were screened every 2-3 days for revertants, i.e., animals that lost the twitching phenotype.

### Small RNA sequencing and bioinformatic analysis

We collected *C. elegans* samples of wild-type, in triplicate, and 1x each *lotr-1* allele (1x *xf58*, 1x *xf60*, and 1x *xf61*). We subsequently treated these three *lotr-1* allele samples as triplicates. Worms were synchronized by bleaching and overnight hatching in M9. L1 arrested worms were brought on OP50-seeded NGM plates. After 63 hours, gravid adult animals were washed off plates into fresh tubes with M9 buffer and frozen in dry ice. For RNA extraction, worms were thawed and M9 was replaced with 250 µL of worm lysis buffer (0.2M NaCl, 0.1M Tris pH 8.5, 50 mM EDTA, 0.5% SDS), supplemented with 300 µg Proteinase K (Sigma-Aldrich, P2308). Subsequently, lysis was conducted for approximately 90 minutes at 65°C. Next, 3 volumes of TRIzol LS (Life Technologies, 10296-028) were added and subsequent isolation was as defined by the manufacturer’s instructions. Samples were enriched for small RNAs using a mirVana Kit (Life Technologies, AM1561).

Samples were subsequently treated with RppH as described.[117] Then, 1 µg of RppH-treated RNA was loaded on a 15% TBE-urea gel and size-selected between 16-to 30-nt. Purified fraction was confirmed by Bioanalyzer sRNA chip (Agilent). Library preparation was based on the NEBNext Multiplex sRNA Library Prep Set for Illumina (New England BioLabs) with slight modifications. To counteract ligation biases, the 3’ and 5’ adapters contained four random bases at the 5’ and 3’-end, respectively, and were chemically synthesized by BioScientific. Adapter-ligated RNA was reverse-transcribed and PCR-amplified for 14 cycles using index primers. The PCR-amplified cDNA construct was purified using AMPure XP beads (Beckman Coulter). The purified PCR reaction was checked on the Bioanalyzer using High Sensitivity DNA chip (Agilent). Size selection of the sRNA library was done on LabChip XT instrument (Perkin Elmer) using DNA 300 assay kit. Only the fraction containing 140-165 bp was pooled in equal molar ratio. The resulting 10 nM pool was denatured to 10 pmol with 5% PhiX spike-in and sequenced as single-read on HiSeq 2500 (Illumina) in rapid mode for 51 cycles (plus 7 cycles index read) using on-board cluster generation.

The raw sequence reads in FastQ format were cleaned from adapter sequences and size-selected for 18-30 base-long inserts (plus 8 random adapter bases) using cutadapt v.2.4 (http://cutadapt.readthedocs.org) with parameters ‘-a AGATCGGAAGAGCACACGTCT -m 26 -M 38’ followed by quality checks with FastQC (http://www.bioinformatics.babraham.ac.uk/projects/fastqc). Read alignment to the *C. elegans* genome (Ensembl WBcel235/ce11 assembly) with concomitant trimming of the 8 random bases was performed using Bowtie v.1.2.2 (http://bowtie-bio.sourceforge.net) with parameters ‘-v 1 -M 1 -y --best --strata --trim5 4 --trim3 4 –S’ and the SAM alignment files were converted into sorted BAM files using Samtools v.1.9 (http://www.htslib.org). WBcel235/ce11 gene annotation in GTF format was downloaded from Ensembl release 96 (ftp://ftp.ensembl.org/pub/) and annotation for transposable elements (LINE, SINE, LTR, DNA and RC) in GTF format was downloaded from the UCSC Table Browser (http://genome-euro.ucsc.edu/cgi-bin/hgTables) RepeatMasker track and merged with the gene annotation GTF file. Aligned reads were assigned to individual small RNA loci and classes using Samtools, GNU Awk, Bedtools v.2.27.1 (http://bedtools.readthedocs.io) and Subread featureCounts v.1.6.2 (http://bioinf.wehi.edu.au/featureCounts/) based on the following criteria: structural reads map sense to rRNA, tRNA, snRNA and snoRNA loci; miRNA reads are between 21 and 24 bases and map sense to mature miRNA loci; 21U-RNA reads are 21 base long, start with T and map sense to piRNA (21ur) loci; 22G-/26G-RNA reads are 22/26 base long, start with G and map antisense to transposons or to exons of protein-coding genes, lincRNAs and pseudogenes. Locus-level differential expression analyses of the small RNA classes (miRNA, 21U-RNA, 22G-RNA and 26G-RNA) were carried out with DESeq2 v.1.20.0 (https://bioconductor.org/packages/release/bioc/html/DESeq2.html) using a significance cutoff of 10% FDR and 2-fold change. Normalized read counts from DESeq2 (S3 Table) were used in the Violin and MA plots. Normalized 22G-RNA coverage tracks in bigwig format were produced using Bedtools and bedGraphToBigWig (http://hgdownload.soe.ucsc.edu/admin/exe/linux.x86_64), and were normalized per million non-structural 18-30 base long reads in each sample. Metagene plots of 22G-RNA read coverage over selected gene sets were prepared using a previously described custom normalization strategy [70], which aims to prevent bias for genes with high 22G-RNA levels. First, genes reported as CSR-1 [118], WAGO [119], or mutator [68] targets were extracted from the GTF gene annotation from Ensembl and 22G-RNA coverage for each gene-set was computed using deepTools v.3.1.0 (https://deeptools.readthedocs.io) with parameters ‘computeMatrix scale-regions --metagene --transcriptID gene --transcript_id_designator gene_id --missingDataAsZero --outFileNameMatrix’. Then, using R v.3.5.1 the resulting count matrix of every sample was cleaned from all-zero rows, 22G-RNA counts for each row(gene) were individually normalized using ‘rowSums’ and cumulative profiles computed by ‘colSums’ were plotted as representative metagene plots. WormMine database (http://intermine.wormbase.org/tools/wormmine/begin.do) was utilized for gene identifier conversion. Sequencing data have been deposited to the NCBI Gene Expression Omnibus (GEO) under accession number GSE172070 [temporary token: ejojuoweztqldsz].

### α**-FLAG Immunoprecipitation**

Immunoprecipitation and mass spectrometry were performed largely as described.[84] Animals were grown either in normal OP50-seeded NGM plates or on OP50 high-density plates. The protocol for the production of the latter was adapted from (Schweinsberg and Grant, 2013, https://www.ncbi.nlm.nih.gov/books/NBK179228/). In short, commercially available chicken eggs were cracked, the yolks isolated and thoroughly mixed with LB medium (50 mL per egg yolk). Then, the mix is incubated at 65°C for 2-3 hours. After cooling down, pre-grown OP50 liquid culture is added to the mix (10 mL per egg). 10 mL of this preparation is poured into each 9 cm plate and plates are decanted the next day. After 2-3 days of further bacterial growth and drying, plates should be stored at 4°C.

Worms expressing 3xFLAG-tagged LOTR-1 and ZNFX-1 were grown and synchronized by bleaching and overnight hatching in M9. Synchronized young adults with no visible embryos were collected 51-56 hours post-plating, washed in M9, followed by a last wash in water, and frozen in dry ice. Prior to IP, samples were thawed, 2x Lysis buffer was added (50 mM Tris/HCl pH 7.5, 300 mM NaCl, 3 mM MgCl_2_, 2 mM DTT, 0.2 % Triton X-100, 1 complete Mini protease inhibitor tablet, #11836153001) and lysis was conducted with sonication in a Bioruptor Plus (Diagenode, on high level, 10 cycles of 30 seconds on/off). Embryos were collected by bleaching gravid animals approximately 72 hours after plating, washing with M9, perform one last wash step in 1x lysis buffer (25 mM Tris/HCl pH 7.5, 150 mM NaCl, 1.5 mM MgCl_2_, 1 mM DTT, 0.1 % Triton X-100, 1 complete Mini protease inhibitor tablet, #11836153001) and by freezing worm pellets in liquid N_2_ in a pre-cooled mortar. Prior to IP, the pellets were ground to a fine powder in a pre-cooled mortar, then transferred to a cold glass douncer and sheared for 40 strokes with piston B. First round of IPs to FLAG-tagged LOTR-1 were performed using 1.5 mg of total embryo or young adult extracts, while for the second round of these experiments 1 mg of embryo or young adult extracts were used. IPs to FLAG-tagged ZNFX-1 were performed using 1 mg of young adult extract and 0.65 mg of embryo extract. IPs were performed in quadruplicates, with exception of FLAG IPs to LOTR-1(ΔTudor) in embryos, which were conducted in triplicate. FLAG-tag immunoprecipitation was performed using an α-FLAG antibody (Monoclonal ANTI-FLAG® M2 antibody produced in mouse, Sigma Aldrich, #F3165) and Protein G magnetic beads (Invitrogen™ Dynabeads™ Protein G; #10004D). 30 µL of beads were used per IP. Beads were washed three times with 1 mL of Wash Buffer (25mM Tris/HCl pH 7.5, 300 mM NaCl, 1.5 mM MgCl_2_, 1 mM DTT, 1 complete Mini protease inhibitor tablet, #11836153001). 2 µg of α-FLAG antibody were incubated with the beads and the extract for approximately 3 hours, rotating at 4°C. Afterwards, samples were washed 5 times with 1 mL of Wash Buffer. Finally, the beads were resuspended in 1x LDS buffer (Thermo Scientific, #NP0007) supplemented with 100 mM DTT and boiled at 95°C for 10 minutes.

### Mass spectrometry

IP samples were boiled at 70°C for 10 minutes and separated on a 4-12% gradient Bis-Tris gel (Thermo Fisher Scientific, #NP0321) in 1x MOPS (Thermo Fisher Scientific, #NP0001) at 180 V for 10 minutes. Then, samples were processed separately, first by in-gel digestion [120,121], followed by desalting with a C18 StageTip.[122] Afterwards, the digested peptides were separated on a heated 50□cm reverse□phase capillary (75 μM inner diameter) packed with Reprosil C18 material (Dr. Maisch GmbH). Peptides were eluted along a 90 min gradient from 6 to 40% Buffer B (see StageTip purification) with the EASY□nLC 1,200 system (Thermo Fisher Scientific). Measurement was done on an Orbitrap Exploris 480 mass spectrometer (Thermo Fisher Scientific) operated with a Top15 data□dependent MS/MS acquisition method per full scan. All raw files were processed with MaxQuant (version 1.5.2.8) and peptides were matched to the *C. elegans* Wormbase protein database (version WS269). Raw data and detailed MaxQuant settings can be retrieved from the parameter files uploaded to the ProteomeXchange Consortium via the PRIDE repository: identifier PXD025186. Reviewer access credentials: Username; Password: pHYaTtrF.

## Acknowledgments

We thank Frederic Bonnet in the MDIBL Imaging Core for assistance with acquiring super-resolution images and Chris Smith in the MDIBL Genome Services Core for sequencing services. We are grateful to Sabrina Dietz and Marion Scheibe for data management and technical assistance. We thank the IMB Genomics core facility and the IMB Media Lab for support in library preparation and providing consumables, respectively. We also thank Michelle Gutwein and Karin Kiontke for assistance with generating transgenic CRISPR strains and Olivia Bay for assistance with imaging. Some strains were provided by the CGC, which is funded by NIH Office of Research Infrastructure Programs (P40 OD010440).

E.A.M was supported by NIH-NIGMS (F32 GM128248). C.S.S. and D.L.U. area supported by NIH-NIGMS (R01 GM113933) with use of equipment supported by NIH-NIGMS (P20 GM103423). Work in R.F.K.’s lab was supported by grants KE1888/1-1 and KE1888/1-2 from the Deutsche Forschungsgemeinschaft. Work in the K.C.G. and F.P. labs was supported by a grant to the NYUAD Center for Genomics and System Biology from the NYUAD Research Institute (ADHPG CGSB) and other funding from NYU Abu Dhabi.

## Supporting information

**Figure S1.**
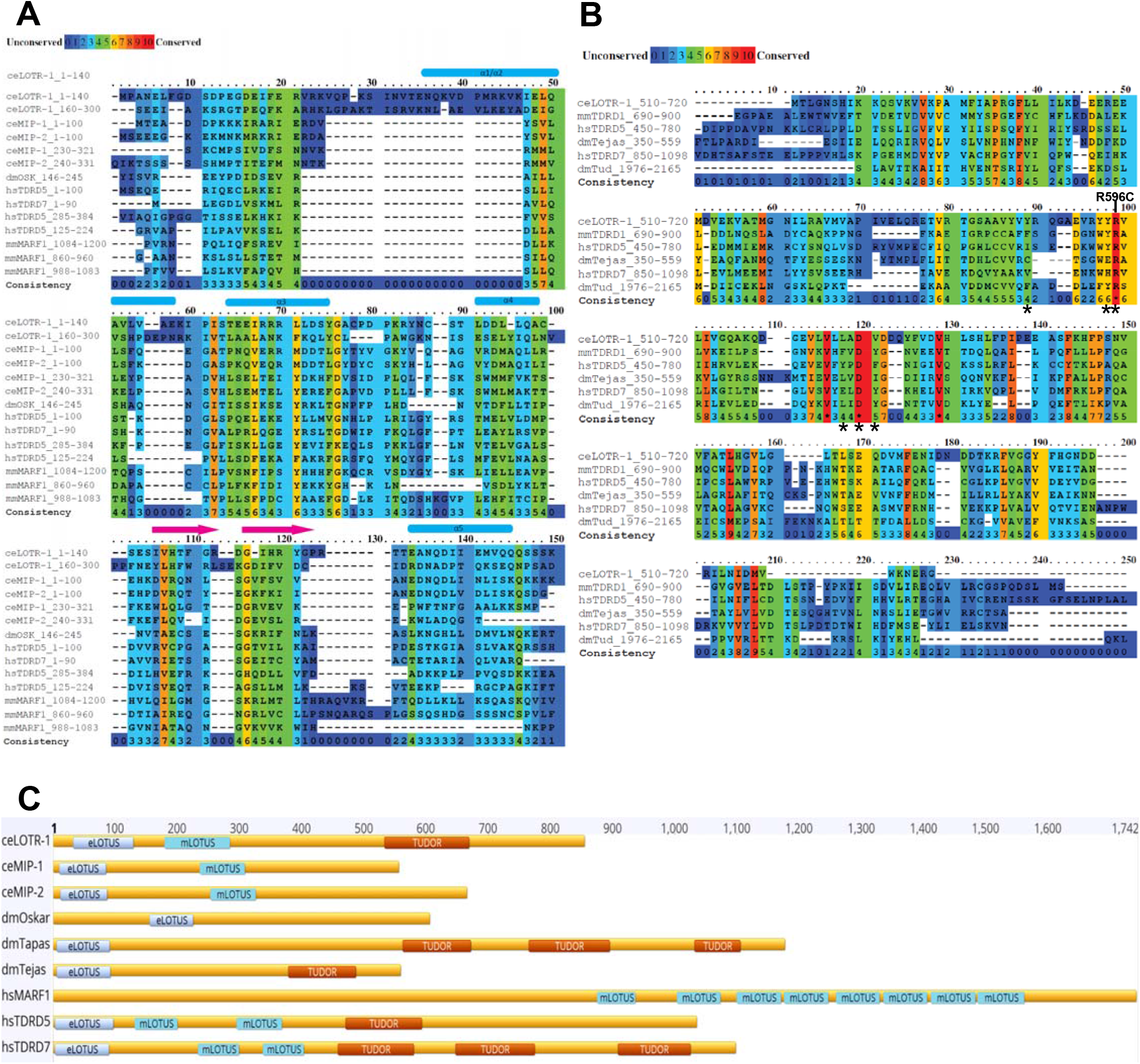
Sequence conservation of LOTUS and Tudor domains across species. Sequence alignment across indicated species for the A) LOTUS domains and B) Tudor domains. C) Conservation of LOTUS and Tudor domains across indicated proteins and species.

**Figure S2.**
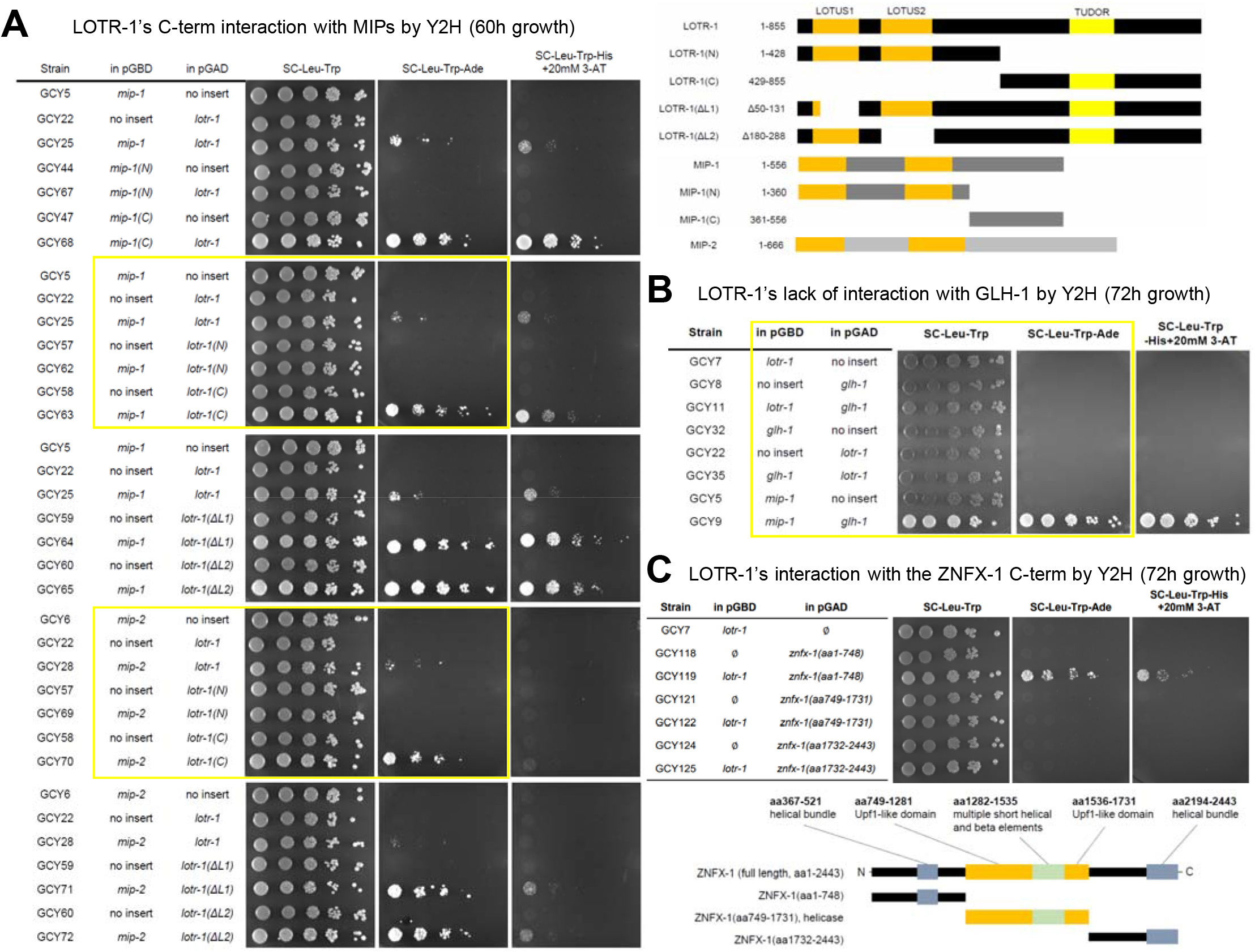
LOTR-1 associates with the MIP proteins and ZNFX-1, but not GLH-1 by Y2H. A) Y2H analysis of MIP-1, MIP-2, and LOTR-1. MIP-1’s C-terminal half interacts with full length LOTR-1. LOTR-1’s C-terminal half interacts with both MIP-1 and MIP-2, and the LOTR-1/MIP interactions are independent of LOTR-1’s LOTUS domains. B) Y2H analysis of LOTR-1, MIP-1 and GLH-1 do not uncover an interaction between LOTR-1 and GLH-1. C) Y2H activation through the N-terminal third of ZNFX-1 is strengthened by an association with LOTR-1. Yellow boxes are duplicated in Fig 4.

**Figure S3.**
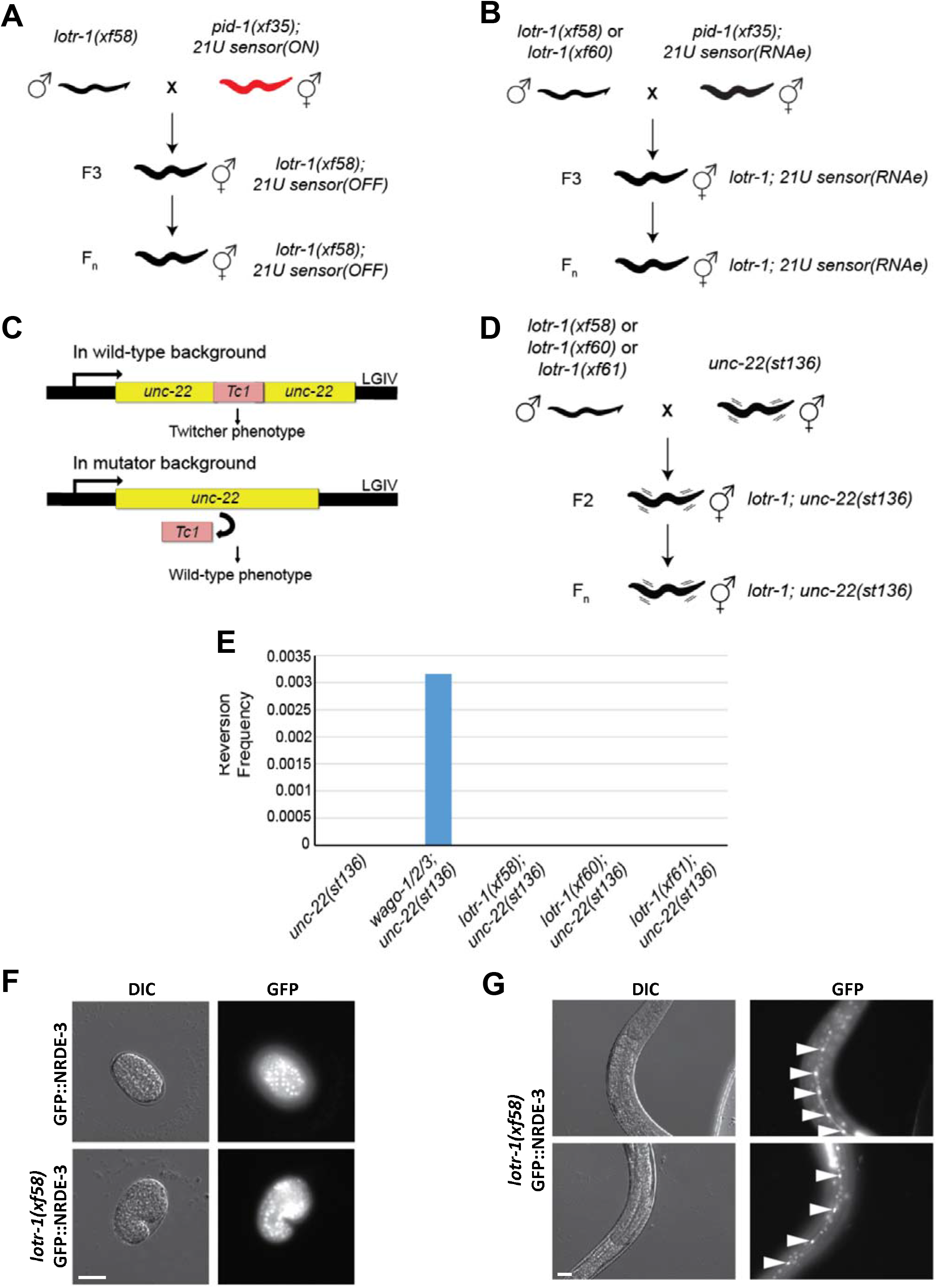
The effect of *lotr-1* mutations of small RNA silencing and transposon mobilization. A) Cross scheme of a non-stably silenced 21U sensor with *lotr-1(xf58)* mutants. The F3 of the indicated genotype was scored for mCherry expression in the germline. B) Schematics of two independent crosses between two *lotr-1* mutant alleles and a 21U sensor that is stably silenced under RNAe. C) Schematic of the *unc-22(st136)* allele. In an otherwise wild-type background, the Tc1 copy integrated in the *unc-22* gene does not mobilize, and these mutants display a twitcher phenotype. However, if transposon silencing is compromised Tc1 will become mobile and transpose leaving an intact *unc-22* gene, which restores the wild-type phenotype. D) Layout of the *unc-22(st136)* x *lotr-1* crosses to address Tc1 derepression. E) Reversion frequency of the indicated genotypes. F) Transgene imaging of the indicated sensor strain. G) Differential interference contrast, and fluorescence photomicrographs of Embryos (G), and L4 animals (H) of the indicated genotypes. GFP is observed in the nuclei of hypodermic seam cells, indicated by white arrowheads in H. The images are representative of at least 10 embryos or 10 L4 worms.

**Figure S4.**
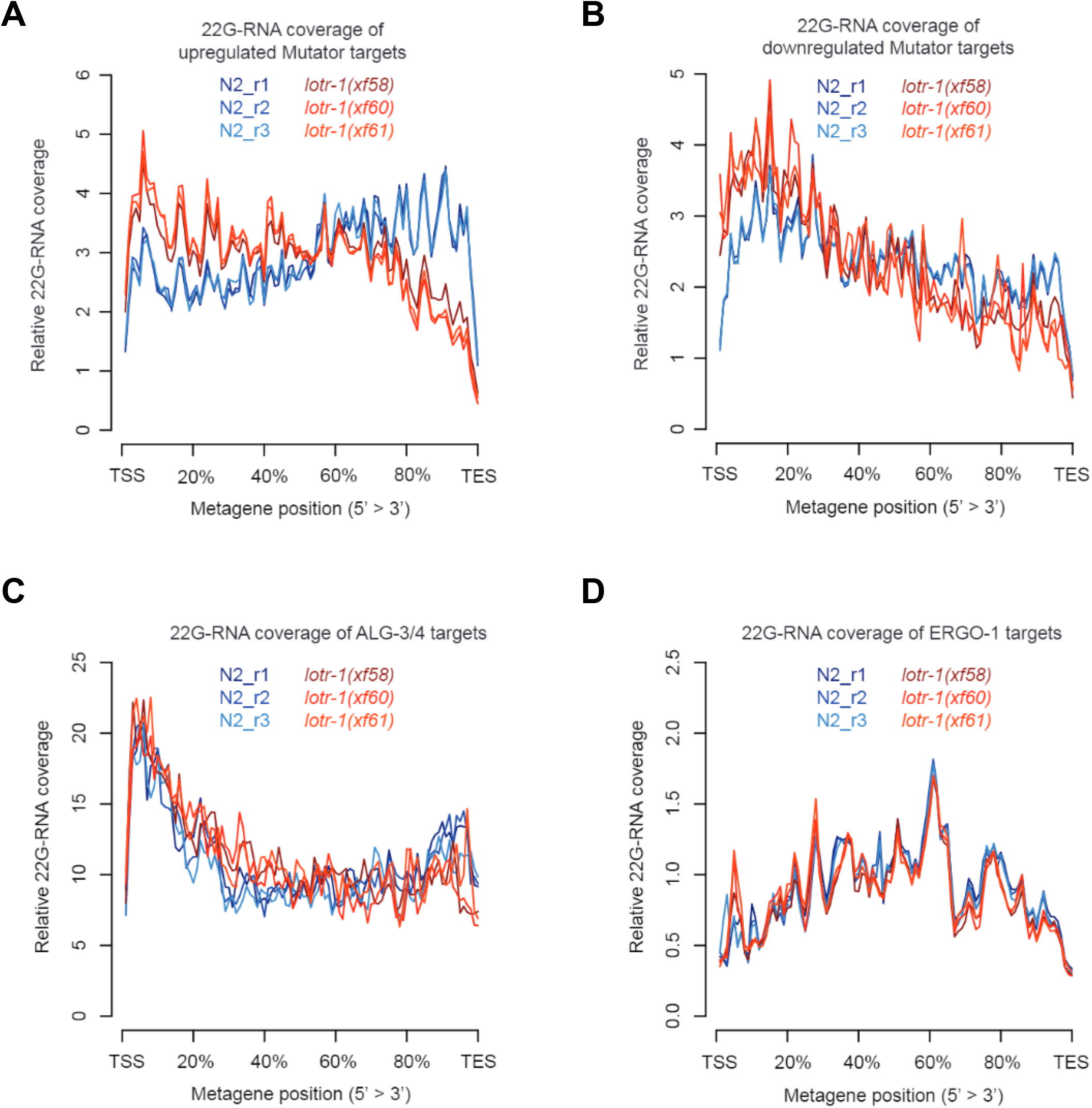
22G-RNA distribution over specific target genes in *lotr-1* mutants. Metagene plots to visualize relative 22G-RNA distribution in wild-type (N2) and *lotr-1* mutants over A) upregulated Mutator targets, B) downregulated Mutator targets, C) ALG-3/4 targets, and D) ERGO-1 targets.

**S1 Table. Enriched Proteins of 3xFLAG::GFP::LOTR-1 anti-FLAG IP-qMS**

**S2 Table. Enriched Proteins of 3xFLAG::GFP::ZNFX-1 anti-FLAG IP-qMS**

**S3 Table. 22G-RNA and 26G-RNA differential expression analysis with gene annotation.**

**S4 Table. Strains used or created for this study**

